# Functional validation of human SK channels variants causing NEDMAB and Zimmermann-Laband syndrome-3 in *C. elegans*

**DOI:** 10.1101/2025.04.11.648345

**Authors:** Sara Sechi, Charlotte Galaup, Maelle Jospin, Thomas Boulin

## Abstract

Small-conductance Ca^2+^-activated K^+^ channels (SK channels) are widely expressed in the central nervous system, where they play a crucial role in modulating neuronal excitability. Recent studies have identified missense variants in the genes encoding SK2 and SK3 channels as the cause of two rare neurodevelopmental disorders: NEDMAB and ZLS3, respectively. Here, we used *C. elegans* as an *in vivo* model to investigate the functional consequences of these patient variants. The *C. elegans* ortholog KCNL-1 regulates neuronal and muscle excitability in the egg-laying system, a well-characterized model circuit. To visualize KCNL-1 expression and localization, we generated a fluorescent translational reporter at the endogenous *kcnl-1* locus. We then introduced eight point mutations corresponding to pathogenic variants reported in NEDMAB or ZLS3 patients. Our study confirmed the molecular pathogenicity of the ZLS3-associated mutations, revealing a gain-of-function effect that led to increased *in utero* egg retention, likely due to electrical silencing of the egg-laying circuitry. NEDMAB mutations exhibited more complex phenotypic effects. Most caused a loss-of-function phenotype, indistinguishable from null mutants, while one displayed a clear gain-of-function effect. Additionally, a subset of NEDMAB variants altered KCNL-1 localization, suggesting an impairment in channel biosynthesis, trafficking or stability. These findings provide new insights into the molecular mechanisms underlying NEDMAB and ZLS3 physiopathology, enhancing our understanding of SK channel dysfunction in human disease. Moreover, they establish *C. elegans* as a robust and cost-effective *in vivo* model for rapid functional validation of new SK channel mutations, paving the way for future investigations.

## Introduction

Small conductance Ca^2+^-activated K^+^ channels (SK channels) are voltage-independent K^+^ channels that are activated by sub-micromolar increases in intracellular Ca^2+^ concentration. Expressed throughout the central nervous system, these channels play an important role in the regulation of membrane excitability. By modulating the afterhyperpolarization duration of action potentials, they regulate the firing frequency of neurons. They also contribute to synaptic plasticity by influencing the induction of long-term potentiation. SK channels are composed of four subunits, each consisting of six transmembrane domains, a pore loop and a calmodulin-binding domain (CaMBD) at the C-terminus. The CaMBD is formed by two alpha-helices (HA and HB) and is constitutively bound to calmodulin (CaM). In humans, four genes, named *KCNN1* to *4*, encode the small (hSK1, hSK2 and hSK3) and intermediate (hSK4) conductance Ca^2+^-activated K^+^ channels (Adelman, Maylie, and Sah 2012; Sun et al. 2020; Rahman et al. 2023).

Dysregulation of SK channel activity has been reported in animal models of several neurological disorders, including schizophrenia, Alzheimer’s and Parkinson’s diseases (Sun et al. 2020). More recently, mutations in *KCNN* genes have been identified in patients suffering from rare congenital neurological disorders (Nam, Downey, et al. 2023). Seven variants in *KCNN3* have been described in association with the Zimmermann-Laband Syndrome 3 (ZLS3, OMIM #618658), which is characterized by mild to moderate developmental delay, intellectual disability, hypotonia, craniofacial dysmorphism, gingival enlargement, hypertrichosis, and nail hypoplasia (Bauer et al. 2019; Gripp et al. 2021; Schwarz et al. 2022). Variants in *KCNN2* are also associated with neurodevelopmental pathologies. They were first identified in two patients mainly suffering of movement disorders (Raghuram et al. 2017; Balint et al. 2020). Later, Mochel and co-workers described eleven patients with impaired intellectual development and movement abnormalities, who carried heterozygous mutations in *KCNN2*. The pathology associated with *KCNN2* mutations was named NEDMAB for NEurodevelopmental Disorder with or without variable Movement or behavioral ABnormalities (OMIM #619725) (Mochel et al. 2020). More recently, a variant in *KCNN2* has been described in a familial form of essential tremor (ET), associated with anxiety and mild to severe cognitive impairment (d’Apolito et al. 2023).

Functional validation is required to establish that variants are causative for disease. The impact of several *KCNN* patient mutations on human SK channel activity was investigated using electrophysiological recordings in heterologous systems (Nam, Downey, et al. 2023; Rahman et al. 2023). Three KCNN3/hSK3 variants, S436C, K269E and G350D, were expressed in CHO cells and resulted in gain-of-function behavior of the mutant hSK3 channel, characterized by faster activation following Ca^2+^ diffusion through the patch-clamp pipette (Bauer et al. 2019). Regarding KCNN2/hSK2 variants, Mochel and co-workers studied the effects of six variants, five point mutations E30Q, I359M, G362S, L388V, L432P, and one single residue deletion, L321del. When expressed in CHO cells, wild-type and E30Q KCNN2/hSK2 channels generated large currents within a few seconds after breakthrough to the whole-cell configuration, whereas no current was detected for I359M, G362S, L388V, L432P and L321del (Mochel et al. 2020). Based on these results, the authors concluded that these last five variants led to a loss-of-function of hSK2 channels. These results were reproduced in HEK293 cells expressing rat KCNN2/SK2 channels, and a loss-of-function phenotype was also observed for a new patient variant, Y361C(Nam, Rahman, et al. 2023). However, in both studies, the authors could not determine whether this loss-of-function phenotype was due to a decrease in channel activity or an alteration in the biosynthesis, trafficking or stability of the protein.

The behavior of the channels expressed in a heterologous system may not reflect that in the native environment. Therefore, functional validation also requires *in vivo* studies. To date, only two patient variants, both in *KCNN2*, have been reproduced in animal models, one in mouse (Raghuram et al. 2017) and one in rat (Kuramoto et al. 2017). Generating variants in mammalian organisms is expensive and time-consuming. It may not be the most appropriate approach to match the increasing number of variants identified in rare diseases, thanks to improvements in next-generation sequencing. The use of non-mammalian animal models, such as the nematode *Caenorhabditis elegans*, has proven to be a successful strategy to rapidly and cost-effectively assess the *in vivo* effects of gene variants, provided the gene is well conserved (Yamamoto et al. 2024).

The *C. elegans* genome contains four genes encoding SK channel subunits. Two of them have been studied previously. *kcnl-1* and *kcnl-2* share a similar expression profile, with expression in head and tail ganglia, the ventral nerve cord, and in the egg-laying system (Chotoo et al. 2013; Vigne et al. 2021). *kcnl-2* has been proposed as a modifier gene in *C. elegans* models of spinal muscular atrophy and amyotrophic lateral sclerosis (Nam et al. 2018; Dimitriadi et al. 2013). *kcnl-2* null mutants exhibit mild phenotypic defects, with a slight reduction of locomotion, pharyngeal activity and egg-laying (Dimitriadi et al. 2013; Chotoo et al. 2013). *kcnl-1* mutations also affect egg-laying: the V530L gain-of-function mutation, found in a natural isolate of *C. elegans*, leads to hypoactivity of the system, with more eggs retained in the uterus, whereas a putative loss-of-function mutation causes hyperactivity of the egg-laying system with fewer eggs retained (Vigne et al. 2021).

In this study, we used CRISPR/*Cas9* gene editing to introduce two human *KCNN3* and six *KCNN2* variants into the *C. elegans* orthologue *kcnl-1*. This gene was chosen because both gain and loss-of-function mutations have clear effects on the egg-laying system (Vigne et al. 2021). Our results showed that ZLS3-hSK3 mutations also result in a gain-of-function effect in *C. elegans*, with a significant egg retention and laying of older embryos. Conversely, four *KCNN2* mutations identified in NEDMAB and related pathologies exhibited a loss-of-function phenotype, characterized by fewer eggs retained *in utero* and premature embryos laid. Strikingly, the I359M NEDMAB mutation caused gain-of-function and not loss-of-function effects. All the variants were engineered in the endogenous *kcnl-1* locus which was modified to encode a version of KCNL-1 tagged at its C-terminus with the fluorescent protein wrmScarlet. Visualization of the fluorescent SK channel revealed that the two ZLS3 variants and three of the six NEDMAB mutants do not show obvious alterations in biosynthesis, trafficking or protein stability, while the remaining three mutants exhibit clear disruptions of their fluorescence pattern. Overall, our results indicate that *C. elegans* is an excellent system to easily assess the functional impact of ZLS3 and NEDMAB mutations.

## Materials and methods

### *C. elegans* genetics and molecular biology

Worms were grown at 20 °C on nematode growth medium (NGM) with *Escherichia coli* OP50 as a nutrient source, according to standard protocols (Brenner 1974).

All the strains used in this study, listed in Supplementary Table ST1, were generated by CRISPR/*Cas9* gene editing, performed according to published protocols (Ben Soussia et al. 2019). CRISPR/*Cas9-*induced molecular lesions were confirmed by Sanger sequencing for all alleles generated in this study. Germline transformation was performed by microinjection into the gonad of one-day old *C. elegans* adult hermaphrodites. All crRNA, single-strand oligonucleotides, repair templates and sequencing oligonucleotides used to generate and validate the different alleles are listed in Supplementary Table ST2.

A fluorescent version of *kcnl-1* was generated by inserting the wrmScarlet fluorescent protein (El Mouridi et al. 2017) at the C-terminus of KCNL-1. The strains carrying the patient variations were then engineered in this genetic background (Supplementary Table ST3). To ensure that the observed phenotype was due to the patient variant and not to off-target mutations, two or three independent alleles were generated for each patient variant.

The *kcnl-1* knock-out strain was generated by deleting the last five exons (from exon 10 to 14 considering isoform c, B0399.1c.1), which encode all the functional domains common to all isoforms of the protein.

### Behavioral assays

Three phenotypic assays were used to assess egg-laying activity: *in utero* egg accumulation, laid-embryos staging, and brood size assays.

For the *in utero* accumulation assay, synchronized L4 larvae were collected from independent plates, and placed at 20°C for 30 hours. The animals were then placed individually in a drop of diluted commercial bleach (1%). After cuticle dissolution (about 10 minutes), the number of eggs was counted. Approximately 30 animals were analyzed per replicate. Data collected from two replicates were combined into superplots (Lord et al. 2020), with each replicate represented by one color.

To determine the stage at which embryos were laid, 30 synchronized adult animals were incubated at 20°C for 30 minutes. The embryos laid during this time were then placed on ice to slow down their development, mounted on a slide and observed with an inverted microscope at 400x magnification. They were scored into one of the following three categories: less than 20 cells, 20 cells to gastrula, post-gastrula.

To measure brood size, late L4 worms were individually placed on NGM plates for 24 hours at 20°C and transferred to a new plate every 24 hours for a total of 72 hours. The plates containing the offspring were incubated overnight at 25°C and the number of larvae on each plate was counted.

### *In vivo* imaging

Worms were immobilized on 2% agarose dry pads mounted in M9 buffer (3 g KH_2_PO_4_, 6 g Na_2_HPO_4_, 5 g NaCl, 0.25 g MgSO_4_·7 H_2_O, and distilled water up to 1 L) containing 2% polystyrene beads (Tebu-Bio, polystyrene polybeads 0.10 μm microsphere). Epifluorescence images were collected from young adult worms, using an AZ100 macroscope (Nikon) equipped with a Flash 4.0 CMOS camera (Hamamatsu Photonics). Confocal imaging was performed under an inverted confocal microscope (Olympus IX83) equipped with a confocal scanner unit spinning-disk scan head (Yokogawa) and an EMCDD camera (iXon Ultra 888, Andor). Head and vulva images were obtained from young adults using 40x and 60x objectives respectively. Both epifluorescence and confocal images were visualized using the open-source image processing software Fiji ImageJ (version 2.3.0/1.53q).

### Statistics

Statistical analyses for embryo accumulation in the uterus and brood size were performed using Prism 10 (version 14.4.1). For embryo accumulation, a superplot was created, while the data pool was subjected to statistical comparison. The Mann-Whitney test was used for the embryo accumulation assay and the brood size assay in ZLS3 mutants, while either Ordinary One-Way ANOVA or Kruskal-Wallis was applied for NEDMAB mutants, depending on the normality of data distributions. The specific statistical tests employed, along with corresponding P-values, are reported in Supplementary Table ST4. Graphical representations were also generated using Prism.

To analyze the distribution of embryo developmental categories, a custom Python script was developed in the Jupyter Notebook environment (Version 7.0.8), with support from ChatGPT. The script, available in “Appendices”, begins with data import and preprocessing to ensure a clean and analyzable dataset. Descriptive statistics were calculated using NumPy and pandas to summarize the main dataset features. Hypothesis testing was performed with SciPy for statistical inferences, specifically using Fisher’s exact test with Bonferroni correction. Data visualization was conducted using Matplotlib and Seaborn, with each raw count for every category normalized to a percentage scale of 100.

## Results

### KCNL-1 is a well-conserved ortholog of human hSK2 and hSK3 potassium channels

Four genes encode SK channel subunits in the *C. elegans* genome: *kcnl-1, kcnl-2, kcnl-3, and kcnl-4*. Overall sequence conservation within core protein domains (i.e., from transmembrane domain S1 to the calmodulin binding domain CaMBD) between KCNL-1 and hSK2 or hSK3 was 43% identity and 68% homology (Fig. 1A). Sequence conservation was notably higher in specific regions that included human variants (Fig. 1A, 1B). Notably, sequence identity was 67% in the S_45_A domain, 100% in the selectivity filter, 41% in the S6 transmembrane segment, and 61% in the calmodulin-binding HA helix (Fig. 1A). Specifically, out of nine pathogenic mutations, seven affected residues that were strictly conserved in KCNL-1, and two variants altered residues that were homologous (Ala instead of Leu for SK2 L388V; Phe instead of Leu for SK2 L432P).

**Figure 1.**
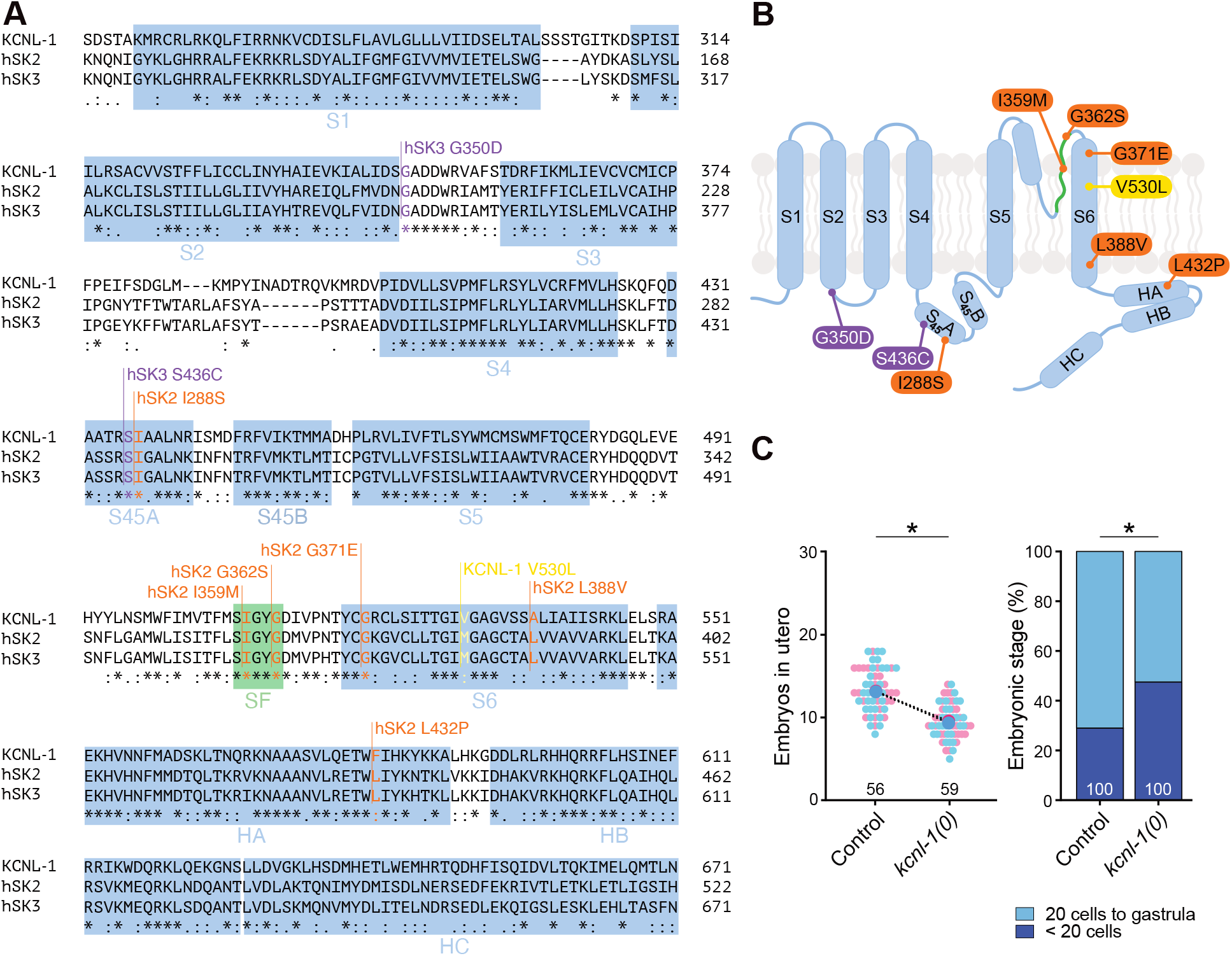
The KCNL-1 channel, a well-conserved ortholog of human SK2 and SK3, regulates egg-laying behavior in *C. elegans*. A. Amino acid sequence alignment of the core domain of the *C. elegans* KCNL-1 channel subunit with human SK2 and SK3 channel subunits. All alpha-helices are labeled in blue. S1–S6 are transmembrane segments, S_45_A and S_45_B represent the two helices in the S4-S5 linker. HA, HB, and HC correspond to helices A, B, and C, respectively. SF, in green, shows the selectivity filter. Patient mutations are marked with purple lines for ZLS3-hSK3 variants, orange lines for NEDMAB-hSK2 variants, and a yellow line for the mutation found in natural isolates. B. Schematic representation of one SK subunit showing 6 transmembrane segments (S1-S6), two alpha-helices S_45_A and S_45_B in the S4-S5 linker, and three C-terminal alpha helices HA, HB and HC. Purple capsules indicate ZLS3-hSK3 patient variants, orange capsules NEDMAB-hSK2 variants, a yellow capsule the native gain-of-function mutant in *kcnl-1*. C. *Left panel, in utero* egg accumulation. Small pink and blue dots represent individual animals from two replicates. Large dots indicate the mean. Mann-Whitney test, * p < 0.05. *Right panel*, distribution of developmental stages of embryos laid over 30 min by 30 animals. Fisher’s exact test with Bonferroni correction, * p < 0.05.

### The SK channel KCNL-1 regulates egg-laying activity in *C. elegans*

In self-fertilizing *C. elegans* hermaphrodites, embryos are retained in the uterus before being expelled through the vulva around the 20-cell stage. We previously showed that the SK channel ortholog KCNL-1 is expressed in egg-laying muscles and neurons using a transcriptional reporter knock-in line (Vigne et al. 2021). To be able to model and analyze the impact of human variants on the biosynthesis and trafficking of KCNL-1 proteins *in vivo*, we engineered a translational knock-in line that labels all KCNL-1 isoforms with the red fluorescent protein wrmScarlet (Fig. S1A). The expression profile of this knock-in reporter was consistent with the transcriptional expression pattern of *kcnl-1*. It showed expression in a subset of head and tail neurons, in addition to clear expression in neurons and muscles of the egg-laying apparatus (Fig. S1B).

We showed previously that KCNL-1 gain-of-function mutations found in natural isolates cause strong embryo retention, and further demonstrated the opposite effect of *kcnl-1* inactivation using a putative reduction-of-function allele (Vigne et al. 2021). Here we used CRISPR/*Cas9* gene editing to generate a large deletion allele that removed all exons encoding key functional domains of KCNL-1 (Fig. S1A), and we used the KCNL-1-wrmScarlet knock-in line as a starting strain to ensure consistency of genetic backgrounds in subsequent experiments. We could show that this molecular null allele of *kcnl-1* increased activity of the egg-laying system using two functional assays. First, we monitored the accumulation of embryos in the uterus at a specific time point (L4 + 30 h at 20°C). Contrary to gain-of-function mutants, *kcnl-1* null animals retained significantly fewer embryos than control animals (Fig. 1C).

Second, we determined the developmental stage of freshly laid embryos. The age of freshly laid embryos is a direct readout of the time spent by an embryo *in utero*. In *kcnl-1* null mutants, we observed a clear change in the distribution of embryo ages. Indeed, significantly more embryos were laid before the 20-cell stage in the absence of KCNL-1 than in control (Fig. 1C). Taken together, these data demonstrate that the SK channel KCNL-1 contributes to the regulation of the egg-laying activity of *C. elegans*.

### ZLS3-hSK3 gain-of function mutations increase KCNL-1 activity and impair egg-laying in *C. elegans*

Naturally occurring gain-of-function mutations – *e*.*g*., KCNL-1 V530L – cause a strong reduction of egg-laying activity, leading to excessive embryo retention and internal matricidal hatching (Vigne et al. 2021). We therefore expected a similar phenotype for gain-of-function mutations associated with ZLS3. We used CRISPR/*Cas9* gene editing to engineer the G350D and S436C patient variants at orthologous positions in the KCNL-1-wrmScarlet control background (Fig. 1A, 2A, Supp. Tab. ST3). In addition, we reproduced the V530L natural variant as a reference allele for hyperactivation of KCNL-1.

ZLS3 knock-in variants could be readily identified during the gene editing procedure, as they produced animals with visible accumulation of embryos *in utero* (Fig. 2B). To determine the impact of ZLS3 mutations, we first quantified the accumulation of embryos *in utero*. While S436C and V530L mutants showed an approximative 2-fold increase in embryo retention (Fig. 2B, 2C, S2A), the effect of G350D was milder (Fig. 2B, 2C). Furthermore, increased *in utero* egg retention was not significantly different from control in two additional independent alleles generated by CRISPR/*Cas9* gene editing (Fig. S2B), indicating that this assay may not be sensitive enough for mutations with a small effect.

**Figure 2.**
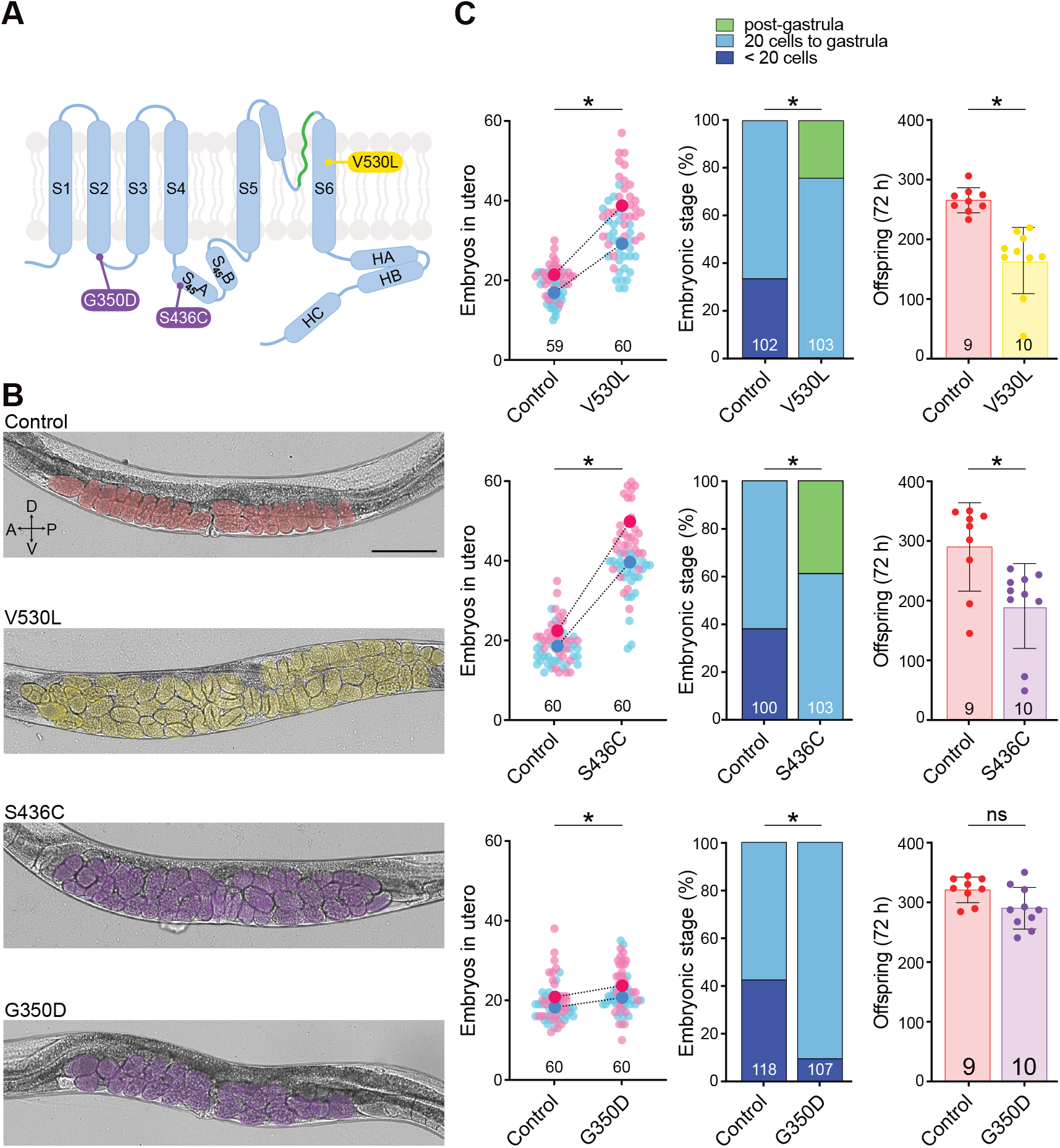
ZLS3-hSK3 mutations impair egg-laying in *C. elegans*. A. Schematic representation of one SK subunit showing the positions of ZLS3-hSK3 S436C and G350D mutations in purple, and the V530L natural variant in yellow. B. Representative brightfield images of L4 + 30 h old worms in a lateral view. Embryos are highlighted in pseudo colors. Number of eggs *in utero*: Control, 18; V530L, 54; S436C, 39; G350D, 20. Scale bar, 200 μm. C. *Left panel, in utero* egg accumulation. Small pink and blue dots represent individual animals from two replicates. Large dots indicate the mean. Mann-Whitney test, * p < 0.05. *Middle panel*, distribution of developmental stages of embryos laid over 30 min by 30 animals. Fisher’s exact test with Bonferroni correction, * p < 0.05. *Right panel*, total brood size over 72 hours. Mann-Whitney test, * p < 0.05.

Next, to support these observations, we measured the developmental stage of freshly laid embryos. Consistent with our expectations, S436C and V530L mutants laid “older” embryos, *i*.*e*., at a more advanced developmental stage (“post-gastrula”) than the control strain (Fig. 2C). Furthermore, this assay also confirmed the mild gain-of-function effect of G350D, since two out of three knock-in lines also laid significantly fewer early stage (<20 cells) and more late-stage embryos (20 cells to gastrula), but no post-gastrula embryos (Fig. S2B). This demonstrates that G350D has a gain-of-function effect in *C. elegans*, albeit with a milder phenotype than S436C and V530L.

Lastly, we performed a complementary assay to measure the total brood size of these mutants. As for V530L mutants (Vigne et al. 2021), stronger *in utero* retention in S436C lead to maternal death and a significant reduction in total brood size measured after 72 hours (Fig. 2C). Consistent with its milder impact in the first two functional assays, G350D mutant showed no change in overall brood size, although some instances of maternal death could be observed, which was never the case for control animals.

Taken together, these data show that ZLS3 patient variants also increase SK channel activity *in vivo* in *C. elegans*, as demonstrated for human hSK3 channels using electrophysiological techniques in heterologous expression systems. They illustrate how simple *in vivo* assays measuring the activity of the egg-laying system (i.e., *in utero* egg accumulation and developmental staging) are sensitive readouts that provide functional evidence for the pathogenicity of patient mutations.

### Most NEDMAB-hSK2 variants strongly reduce KCNL-1 activity and phenocopy a molecular null allele

*KCNN2*/hSK2 variants have recently been linked to a novel rare neurological disease named NEDMAB (Mochel et al. 2020). Contrary to ZLS3-hSK3 variants, electrophysiological characterization indicated a loss-of-function effect of patient variants in hSK2, as no current could be recorded when variant channels were expressed in cell culture systems (Mochel et al. 2020; Nam, Rahman, et al. 2023). To determine whether NEDMAB variants also resulted in functional reduction or loss of KCNL-1 activity, we used CRISPR/*Cas9* gene editing to engineer five missense variants (G362S, L432P, L388V, I288S, G371E) into the KCNL-1-wrmScarlet control background (Fig. 1A, 3A, Supp. Tab. ST3). We then assayed their functional impact in comparison to *kcnl-1* molecular null mutants using the *in utero* egg accumulation and developmental staging assays.

We first analyzed four variants that had been already studied using electrophysiology. The G362S, L432P and I288S mutants accumulated fewer eggs and laid them at a more premature developmental stage than null mutants (Fig. 3B, S2C). For the L388V variant, the corresponding residue in the wild-type KCNL-1 sequence was not a leucine, but an alanine (Fig. 1A). Nevertheless, the replacement by a valine resulted in a loss-of-function effect that was clearly visible in the delayed stage of freshly laid embryos, and milder in the egg accumulation assay (Fig. 3C). We next characterized one variant, G371E, that had not previously been studied by electrophysiology. This variant showed a pronounced loss-of-function effect in *C. elegans*, closely resembling the null mutant, both for *in utero* retention and in the stage of freshly laid eggs. Finally, in all these mutants, as in the null, the total brood size was not altered (Fig. S3). Taken together, these results demonstrate that the loss-of-function effect of hSK2 variants can be robustly detected *in vivo* by monitoring *C. elegans* egg-laying behaviour.

**Figure 3.**
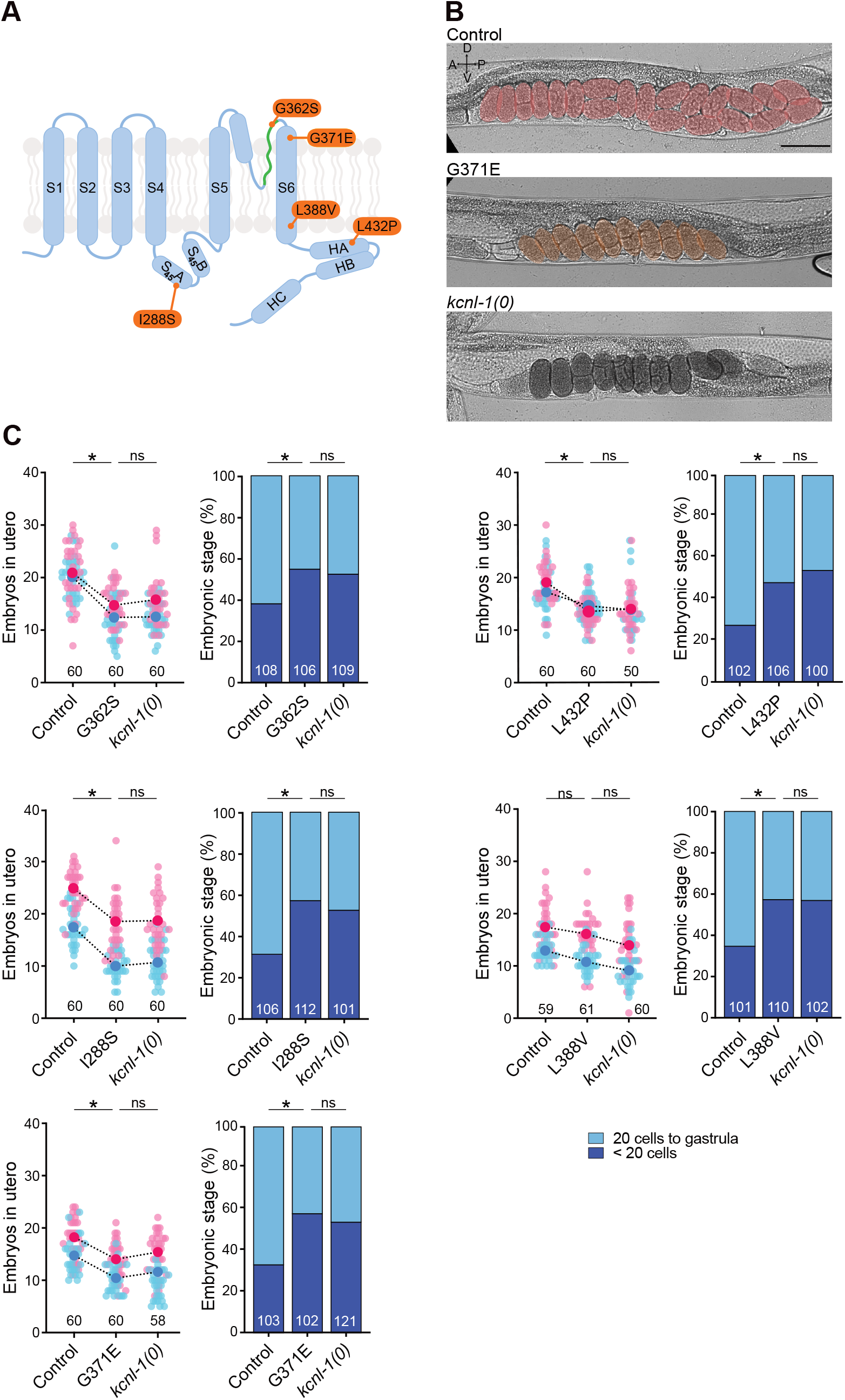
NEDMAB-hSK2 mutations activate the *C. elegans* egg-laying system and phenocopy *kcnl-1* null mutants. A. Schematic representation of one SK subunit with the relative position of five NEDMAB mutations in orange. B. Representative brightfield images of L4 + 30 h old worms in a lateral view. Embryos are highlighted in pseudo colors. Number of eggs *in utero*: Control, 21; G371E, 10; *kcnl-1(0)*, 10. Scale bar, 200 μm. C. *Left panel, in utero* egg accumulation. Small pink and blue dots represent individual animals from two replicates. Large dots indicate the mean. Kruskal-Wallis test, ns p > 0.05, * p < 0.05. *Right panel*, distribution of developmental stages of embryos laid over 30 min by 30 animals. Fisher’s exact test with Bonferroni correction. ns p > 0.05, * p < 0.05.

### The NEDMAB-hSK2 I359M mutation produces a gain-of-function effect in *C. elegans*

The I359M mutant behaved like a loss-of-function channel when studied in heterologous systems (Mochel et al. 2020; Nam, Rahman, et al. 2023). Strikingly, we observed an egg-laying phenotype in *C. elegans* opposite to that of the loss-of-function mutants. Indeed, the I359M mutants displayed a strong *in utero* egg retention, and a high proportion of eggs laid at an advanced developmental stage (Fig. 4B, 4C). In addition, the brood size was reduced due to *in utero* retention and matricidal internal hatching (Fig. 4C). All these phenotypic features were reminiscent of gain-of-function mutant phenotype (Fig. 2C), suggesting that the KCNL-1 I359M, which was described as a loss-of-function variant, actually acts as a gain-of-function mutation *in vivo* in *C. elegans*.

**Figure 4.**
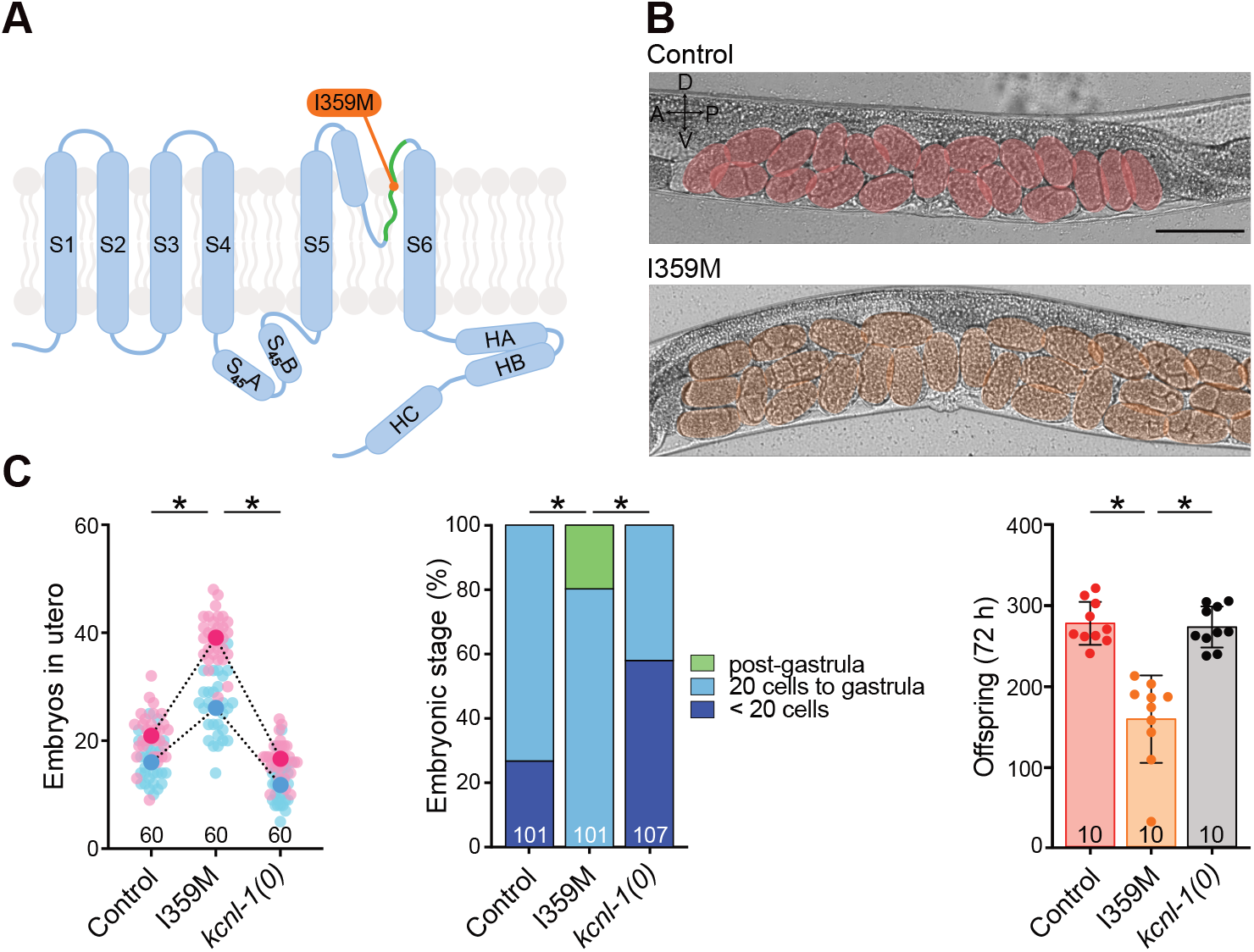
The NEDMAB-hSK2 I359M mutation phenocopies ZLS3 mutations. A. Schematic representation of one SK subunit with the relative position of I359M in orange. B. Representative brightfield images of L4 + 30 h old worms in a lateral view. Embryos are highlighted in pseudo colors. Number of eggs *in utero*: Control, 20; G371E, 30. Scale bar, 200 μm. C. *Left panel, in utero* egg accumulation. Small pink and blue dots represent individual animals from two replicates. Large dots indicate the mean. Ordinary One-Way ANOVA, * p < 0.05. *Middle panel*, distribution of developmental stages of embryos laid over 30 min by 30 animals. Fisher’s exact test with Bonferroni correction analysis. * p < 0.05. *Right panel*, total brood size over 72 hours. Kruskal-Wallis test, * p < 0.05.

### hSK gain-of-function mutations do not alter KCNL-1 localization and protein levels

To investigate the impact of hSK gain-of-function variants on KCNL-1 biosynthesis and subcellular localization, we compared the KCNL-1-wrmScarlet fluorescence pattern in the head of control and gain-of-function mutants. Using an epifluorescence macroscope, we could observe punctate patterns in the somas of pharyngeal and head neurons. Fluorescence was also clearly visible in neurites of the nerve ring and in some projections extending to the tip of the nose (Fig. 5A). Using the same approach, the four gain-of-function mutants (V530L, S436C, G350D, I359M) did not show any gross changes in these KCNL-1 patterns, suggesting that gain-of-function mutations do not dramatically affect the biosynthesis, trafficking or stabilization of the channel (Fig. 5B).

**Figure 5.**
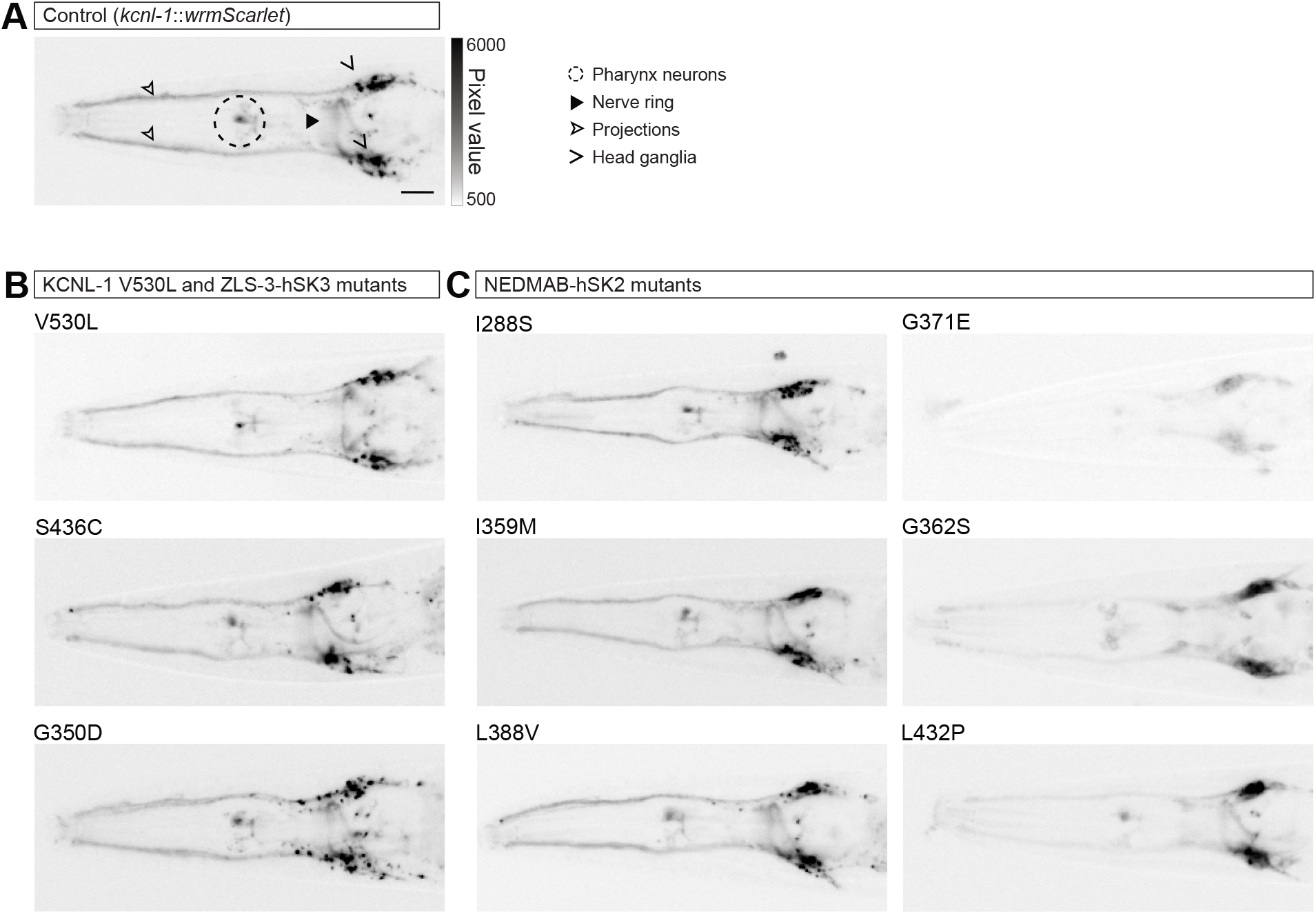
The expression level of KCNL-1 in the head of *C. elegans* is not altered in gain-of-function mutants and is decreased only in a subset of loss-of-function mutants. A. Representative images of the head of control animals. Fluorescence is visible in several neurons of the pharynx (dashed circle), the head ganglia (open arrowheads), projections to the tip of the nose (empty arrowhead) and in the nerve ring (arrowhead). B. Representative images of the head of animals carrying the natural gain-of-function variant V530L or two ZLS3-hSK3 patient variants. C. Representative images of the head of animals carrying NEDMAB-hSK2 patient variants. Images were acquired with an epifluorescence macroscope. Acquisition parameters and dynamic ranges of images are identical between samples. Anterior, left; ventral, down. Scale bar, 20 μm.

### KCNL-1 localization is impaired in a subset of NEDMAB-hSK2 variants

Next, we repeated this analysis to examine the impact of NEDMAB-hSK2 loss-of-function variants on the biosynthesis and subcellular localization of KCNL-1. Two of them, I288S and L388V, did not exhibit gross defects in their fluorescence pattern when compared to control animals (Fig. 5B). In contrast, KCNL-1-wrmScarlet fluorescence was strongly reduced in the three other (G371E, G362S, L432P) mutants (Fig. 5C). To further characterize this decrease, we imaged the head and the vulval region at high magnification using a spinning disc confocal microscope (Fig. 6).

**Figure 6.**
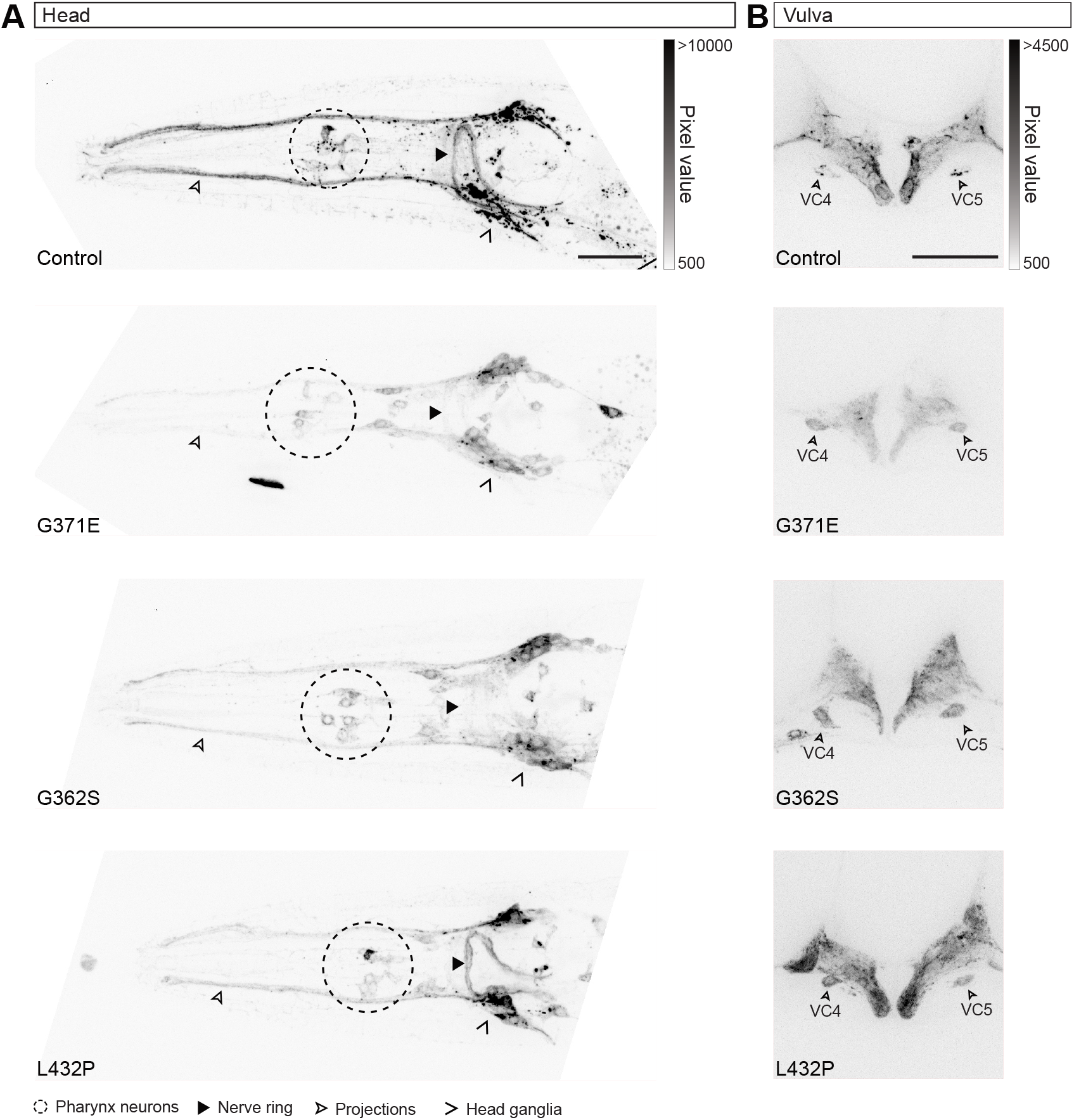
G371E, G362S and L432P NEDMAB-hSK2 variants cause retention of KCNL-1 in the soma of head and vulval neurons. A. Representative images of KCNL-1 localization in the head region. Dorso-ventral views, the neurons in the pharynx are indicated by a dashed circle, the head ganglia by an open arrowhead, projections to the tip of the nose by an empty arrowhead, and the nerve ring by an arrowhead. B. Representative images of KCNL-1 localization in the egg-laying apparatus. Lateral views. Cell bodies of VC4 and VC5 motor neurons are indicated by empty arrowheads. Images were acquired using a spinning disk confocal microscope. Scale bars, 20 μm

We first observed that the fluorescent signal was globally weak in the head of G371E mutants and very faint in the neurites of the nerve ring, as well as in the nose projections. In addition, fluorescence was clearly accumulated in the cell body of pharyngeal neurons and head ganglia, which was not the case in control animals (Fig. 6A, S4). Consistently, the fluorescent signal was also clearly decreased in the egg-laying system (Fig. 6B, S5). We focused our attention on the two VC4 and VC5 neurons that are involved in egg-laying behaviour. In control worms, the outlines of VC4/5 somas were barely visible, because the KCNL-1 signal was enriched in puncta. In contrast, fluorescence was diffuse in VC4 and VC5 in G371E mutants, making the neuronal cell bodies more easily visible. The increased signal in neuronal cell bodies in the head and in the vulva could be due to the retention of the protein in the endoplasmic reticulum, reflecting a defect in channel trafficking.

Next, we examined the G362S mutant, which exhibited similar alterations to the G371E mutant but to a lesser extent. Global fluorescence was decreased, and cell bodies were visible in both the head and in vulva. The signal in the nerve ring and in the nose projections was reduced, but not as much as in G371E (Fig. 6, S4, S5).

Finally, the L432P mutant showed a peculiar KCNL-1 localization in the head: the fluorescence pattern was similar to that of the control in the nerve ring and in a subset of head neurons, while its level was strongly decreased in the projections to the tip of the nose (Fig. 6A, S4). Moreover, some cell bodies were visible in the pharyngeal region. As in G371E and G362S, KCNL-1 fluorescence was decreased overall in the vulva, and the cell bodies of VC4 and VC5 were clearly visible (Fig. 6B, S5).

These results show that the decrease in channel activity observed in NEDMAB loss-of-function mutants could be due to different pathophysiological mechanisms. While all mutations result in a functional reduction of SK channel activity, their effects on channel distribution vary. Some mutations disrupt the normal distribution pattern, whereas others do not produce detectable changes in channel abundance or subcellular localization. This distinction had not been recognized so far.

## Discussion

In this study, we demonstrate that the model nematode *C. elegans* can be effectively used to investigate the functional impact of SK channel variants associated with two rare neurodevelopmental disorders, NEDMAB and ZLS3. Functional validation is an essential step in determining the pathogenicity of variants identified in patients with rare genetic diseases.

Missense mutations pose a specific challenge, as it remains difficult to predict their impact in most cases, and modelling disease mutations, *in vivo*, in animal models represents a gold standard (Yamamoto et al. 2024). To be relevant, a model organism should fulfil three essential requirements. Firstly, its genome should include at least one well-conserved orthologue of the gene mutated in the human pathology. The *C. elegans* genome contains four genes, *kcnl-1, kcnl-2, kcnl-3* and *kcnl-4*, which encode orthologues to human SK channels. Compared to hSK2 and hSK3, the core regions of KCNL-1 and KCNL-2 have the highest percentages of sequence identity (43% and 50%, respectively) and similarity (68% and 69%, respectively). Secondly, engineering mutations in the endogenous loci should be straightforward, time- and cost-efficient. Gene editing in *C. elegans* allows point mutations to be made in as little as three weeks at very low cost (Nance and Frøkjær-Jensen 2019). Finally, the last prerequisite is the ability to assess the effects of mutations by simple and robust functional analysis, allowing many mutations to be tested in parallel. Both KCNL-1 and KCNL-2 have been described to play a role in the well-characterized *C. elegans* egg-laying behaviour (Chotoo et al. 2013; Vigne et al. 2021). We chose to focus our interest on KCNL-1 because gain-of-function and loss-of-function mutations are viable and produce robust phenotypes that are easy to quantify and have little effect on development (Vigne et al. 2021).

Based on electrophysiology results obtained in CHO cells, the ZLS3-hSK3 patient variants were qualified as gain-of-function mutations (Bauer et al. 2019). Indeed, they caused an increase in the apparent Ca^2+^ sensitivity of the mutant channels K269E, G350D and S436C. Here we introduced two of these patient variants, G350D and S436C, at their corresponding position in KCNL-1 (Fig. 1, Supp. Tab. ST3). Both mutations led to the hypoactivation of the egg-laying system, resulting in an increase in the number of eggs retained *in utero* and a developmental shift towards more mature stages of laid embryos. Vigne and co-workers demonstrated strong expression of *kcnl-1* in vulval muscles and VC4/5 motor neurons (Vigne et al. 2021). Transcriptomic data highlighted a high expression of *kcnl-1* in these structures, as well as in HSN motor neurons (Taylor et al. 2021). Egg-laying requires the contraction of vulval muscles, activated by HSN (Emmons 2018). VC4 and VC5 motor neurons also promote egg-laying, directly by acting on the vulval muscles, and indirectly by slowing locomotion and triggering contraction of the striated body-wall muscle cells that aids in egg expulsion (Collins et al. 2016). The phenotypes observed in G350D and S436C mutants are thus consistent with a hyperactivity of KCNL-1. Indeed, increasing K^+^ conductance in all the egg-laying promotive structures should lead to a shutdown of the system.

Previous studies have partially elucidated the mechanism by which G350D and S436C increase the activity of hSK3. G350 and S436 are both located in cytosolic loops. G350 is contained in the S2-S3 linker, while S436 is part of the alpha-helix S45A lying in the S4-S5 linker (Fig. 1). Both residues are located at contact sites between the hSK3 channel and calmodulin in a homology model based on the 3D structure of hSK4 channel (Bauer et al. 2019; Lee and MacKinnon 2018). When expressed in HEK293 cells, the G350D and S436C mutants show an increase in apparent Ca^2+^-sensitivity, which is likely to be at the origin of the gain-of-function effect (Orfali et al. 2022).

In our *C. elegans* model system, the two ZLS3-hSK3 mutations showed a gradation in their gain-of-function phenotypes. S436C channels led to a pronounced inhibition of the egg-laying system, as dramatic as that observed with the V530L mutation found in the natural isolate JU751 (Vigne et al. 2021) and reproduced here in the well-characterized reference isolate N2.

Both exhibited strong retention *in utero*, eggs laid at a stage as advanced as gastrula, and fewer offspring due to matricidal internal hatching. The G350D mutation caused a weaker phenotype: egg retention was moderate, brood size was normal and, although the eggs laid were significantly older compared to control, they never reached the gastrula stage. Interestingly, the apparent Ca^2+^ sensitivity is increased in both mutants compared to wild-type hSK2 (EC_50_ = 0.3 μM), but the extent of this increase is more pronounced in S436C (EC_50_ = 0.087 μM) than in G350D (EC_50_ = 0.19 μM) (Orfali et al. 2022). Therefore, the magnitude of the functional effect of the mutations reflects the phenotypic severity observed in *C. elegans*.

Of the six hSK2 variants identified in NEDMAB and related pathologies, five (G362S, L432P, L388V, I288S, G371E) exhibited a loss-of-function effect similar to that of the molecular null mutant. Three variants (G362S, L388V and L432P) were expressed in heterologous expression systems and did not produce any detectable currents (Mochel et al. 2020; Nam, Rahman, et al. 2023). Loss-of-function mutations can result from modifications of biophysical properties or from changes in biosynthesis, trafficking, localization or stability of the protein. We therefore visualized the impact of mutations on KCNL-1 localization patterns by using an endogenously tagged-version of this SK channel. Among the five loss-of-functions variants, G362S, G371E and L432P show mild to severe defects in the level of KCNL-1 fluorescence level and modification in the distribution pattern, indicative of retention during trafficking. This finding suggests that some NEDMAB mutations may contribute to disease not just by impairing channel function but also by affecting protein localization and stability.

Since the I288S and L388V loss-of-function mutations do not lead to gross changes in the KCNL-1 pattern, they must affect the channel function. The residue I288 is located in the alpha-helix S45A of the S4-S5 linker. In rat SK2, the impact of mutations on the corresponding residue was studied (Kuramoto et al. 2017; Nam et al. 2021; Nam, Rahman, et al. 2023), and the authors concluded that this residue is engaged in a hydrophobic interaction with residue M411 of the HA alpha-helix (Nam et al. 2021). They showed that mutations introducing a less hydrophobic residue destabilize the interaction between the S4-S5 linker of one subunit, the CaMBD of another subunit and the calmodulin. This leads to a less efficient coupling between calmodulin-mediated Ca^2+^ sensing and channel opening through S6 bundle enlargement.

Regarding L388V, it is noteworthy that while the four other loss-of-function mutations exhibit a similar severity of the egg-laying phenotype, the L388V mutant stands out for its milder alterations. Indeed, egg retention was intermediate between wild-type animals and null mutants, and the main change was a premature age of the eggs laid. Interestingly, a mutation near L388, G382D, has been reported in essential tremor, a pathology with less severe clinical symptoms than NEDMAB (d’Apolito et al. 2023). The authors of this study have suggested that G382D and L388V might have a milder effect on hSK2 function and cause a milder clinical picture compared to other variants. Our results support their hypothesis and confirm that our model can be useful to categorize variants based on the severity of their functional effects.

The characterization of I359M, the last NEDMAB-hSK2 mutation we engineered in *C. elegans* KCNL-1, provided unexpected results. While other NEDMAB-hSK2 variants behave as loss-of-function, this mutation caused an unambiguous gain-of-function phenotype. This result is not consistent with the lack of current reported when I359M SK2 channels were expressed in CHO or HEK293 cells (Mochel et al. 2020; Nam, Rahman, et al. 2023). These data should be interpreted with caution, as high variability in SK current intensity has been reported, making it difficult to draw strong conclusions based on current intensity differences alone (Bauer et al. 2019). In addition, channels may behave differently in a heterologous system than in their native context. The I359M mutation is located at the second position of the selectivity filter. The sequence of the filter, SIGYG, is perfectly conserved between KCNL-1 and human SK channels. This sequence is not found in any other mammalian K^+^ channel and is therefore unique to SK channels (Taura et al. 2021). Interestingly, SK2 channels have been reported to exhibit a substantial Na^+^ permeability (Shin et al. 2005). Therefore, I359M could hyperactivate SK2 channels by decreasing this Na^+^ permeability. It remains still puzzling that gain-of-function and loss-of-function mutations could lead to the same symptoms, observed in NEDMAB patients. The I359M mutation requires comprehensive characterization, which falls beyond the scope of this study.

In conclusion, our work establishes *C. elegans* as a new and powerful model system to study the functional effects of SK channel mutations, providing a rapid and cost-effective system for testing the pathogenicity of patient variants. Future research could support this model to screen for potential therapeutic targets for these rare neurodevelopmental disorders. In particular, distinguishing the impact of variants on the biosynthesis versus the function of SK channels in native cellular contexts may be crucial for identifying the most relevant class of therapeutic molecules. This distinction could guide the development of targeted compounds that either enhance proper channel trafficking or restore channel function, depending on the specific defect caused by a given variant.

## Acknowledgments

We thank Sandra Duperrier and Driss Laabid for technical support, Kevin Collins for advice on egg-laying assays. We acknowledge the CIQLE Imaging facility, member of the national infrastructure France-BioImaging supported by the French National Research Agency (ANR-24-INBS-0005 FBI BIOGEN) for access to equipment. This work was performed within the framework of the LABEX CORTEX (ANR-11-LABX-0042) of Université Claude Bernard Lyon 1, within the program “Investissements d’Avenir” (decision n° 2019-ANR-LABX-02) operated by the French National Research Agency (ANR). Strains were generated by SEGiCel (SFR Santé Lyon Est CNRS UAR 3453, Lyon, France) with the support of CNRS and IBiSA. This work was funded by grants to T.B. from Solve-RD (EU Horizon 2020, grant agreement No. 779257) and ANR (AAPG 2020, NBElegAns).

## Data availability

The data that support the findings of this study are openly available in “Appendices”.

**Supplementary Table ST1** List of strains and associated alleles

**Supplementary Table ST2** List of oligonucleotides (crRNA, Repair template and PCR primers) for strain generation and validation

**Supplementary Table ST3** Correspondence between patient variants in hSK2/hSK3 and KCNL-1

**Supplementary Table ST4** Statistics

**Appendix 1** Raw data

**Appendix 2** Python code for statistical analysis of embryonic stage distributions

