## Supplementary material for "Functional validation of human SK channels variants causing NEDMAB and Zimmermann-Laband syndrome-3 in *C. elegans*": SI

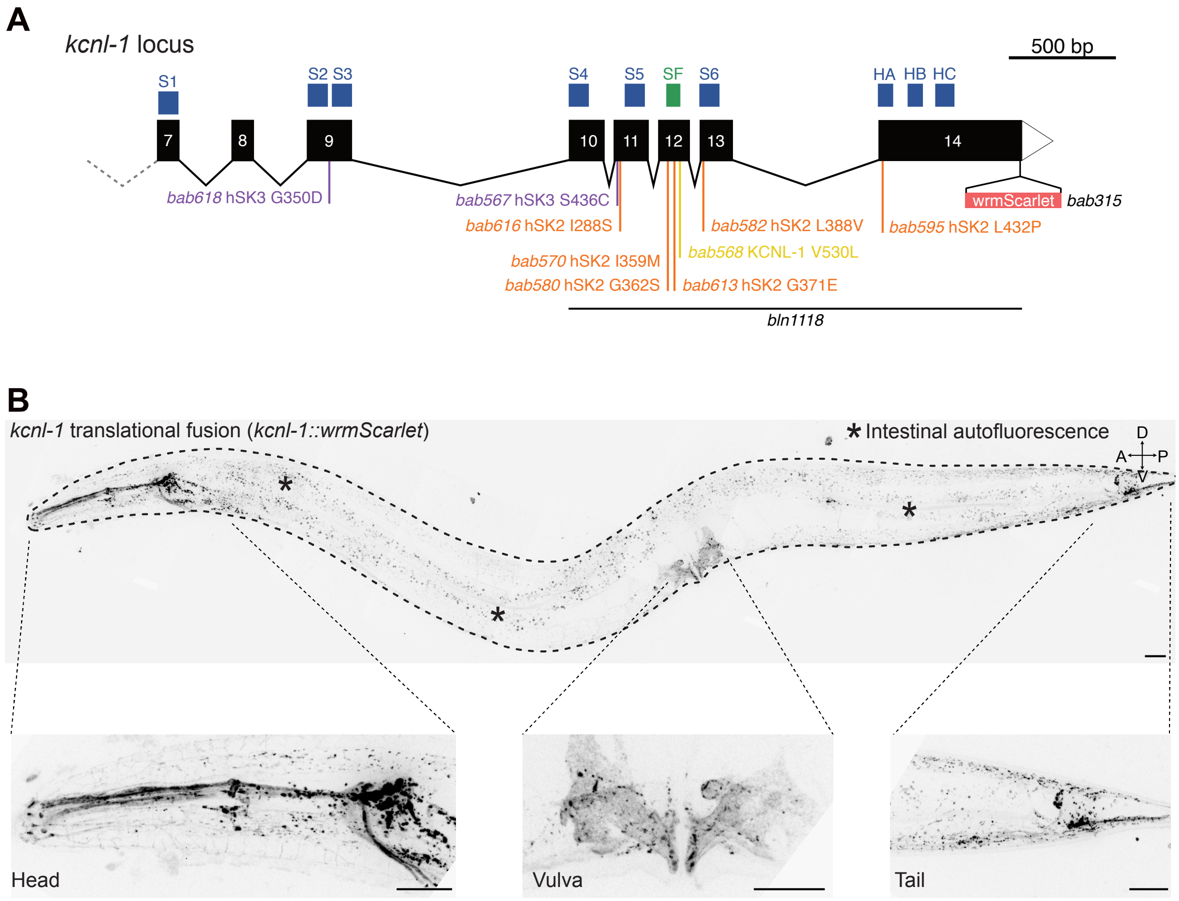

#### Figure S1. *kcnl-1* gene structure and translational reporter.

A. *kcnl-1* gene structure. Common exons (7 to 14) for six isoforms are shown in black. Blue boxes indicate the approximate positions of coding regions for transmembrane segments (S1-S6), the helices A, B and C (HA, HB, HC). The selectivity filter (SF) is labeled in green. The wrmScarlet coding sequence was inserted in frame before the stop codon of the last exon. ZLS3-hSK3 patient variants are indicated in purple, NEDMAB-hSK2 in orange, and the natural gain-of-function mutation in yellow. Alleles are indicated in italics. The extent of the *bln1118* deletion allele is indicated by a black bar.

B. Lateral view of a young adult stage animal for the fluorescent translational knock-in line *kcnl-1(bab315)*. Magnified views of the head, vulval region, and tail. Worm outline, dashed line. Asterisks indicate intestinal autofluorescence. Scale bars, 20 µm.

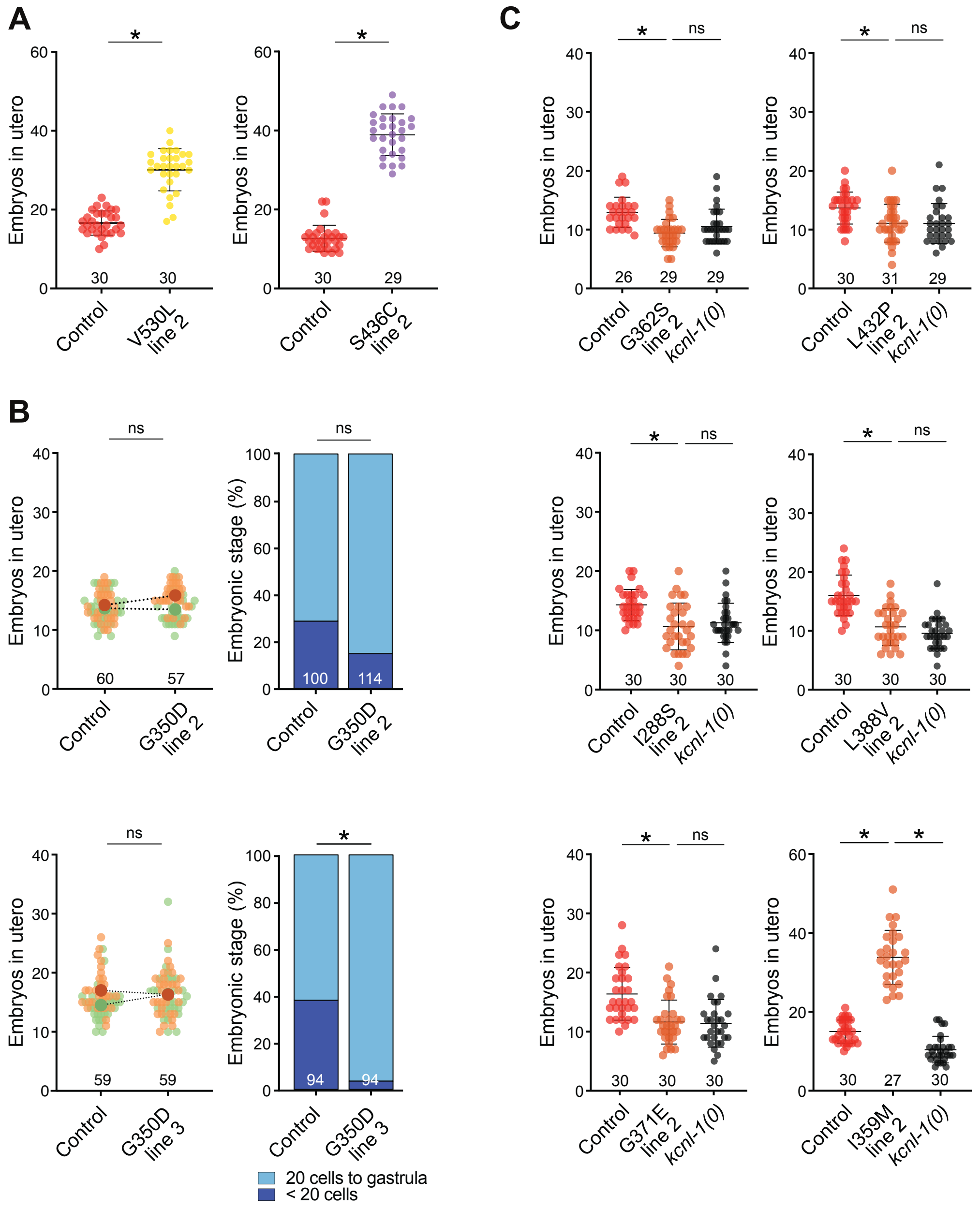

#### Figure S2. Egg-laying phenotypes of additional independent *kcnl-1* alleles.

A. *In utero* egg accumulation for additional independent mutant lines. Mann Whitney, * $p < 0.05$.

B. *Left panel*, *in utero* egg accumulation for two independent G350D lines. Small salmon and green dots represent individual animals coming from two replicates. Large dots indicate the mean. Mann-Whitney test, ns $p > 0.05$.

*Right panel*, distribution of developmental stages of embryos laid over 30 min by 30 animals. Fisher's exact test with Bonferroni correction, ns $p > 0.05$, * $p < 0.05$.

C. *In utero egg accumulation* from independent alleles for NEDMAB-hSK2 variants. Kruskal-Wallis test for I288S, G362S, L388V and L432P, ns $p > 0.05$, * $p < 0.05$. Ordinary One-Way ANOVA for G371E and I359M, ns $p > 0.05$, * $p < 0.05$.

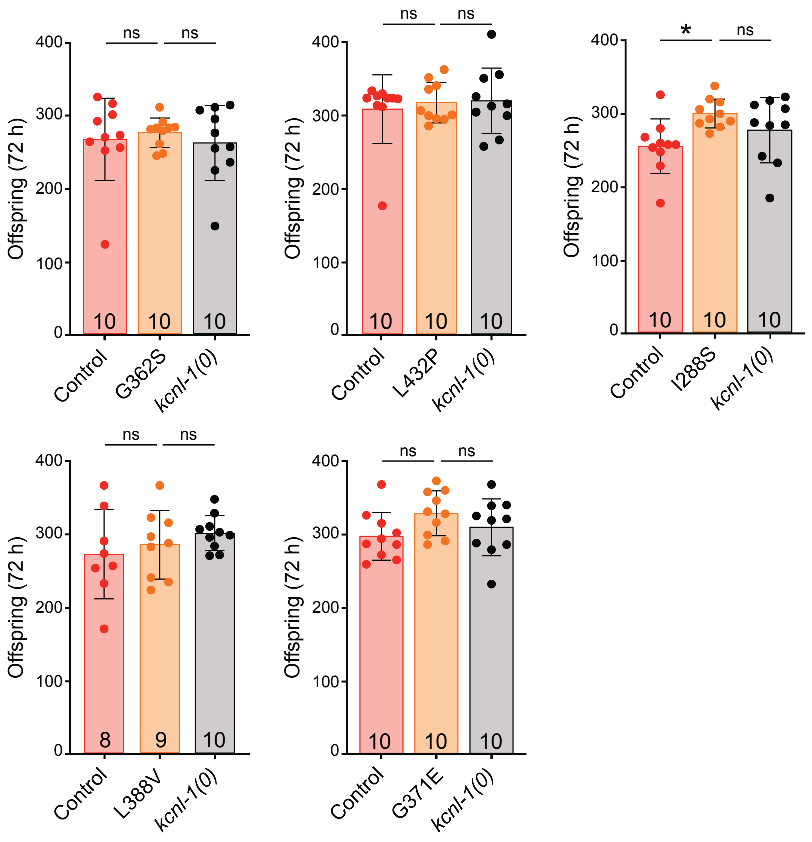

**Figure S3. I288S, G371E, G362S, L432P, L388V and null mutations do not affect brood size.**

Brood size over 72 hours for control animals or mutants carrying the patient variants G362S, L432P, L388V, I228S or G371E. Kruskal-Wallis tests were performed for G362S and L432P, while Ordinary One-Way ANOVA were performed for G371E, I288S and L388V.

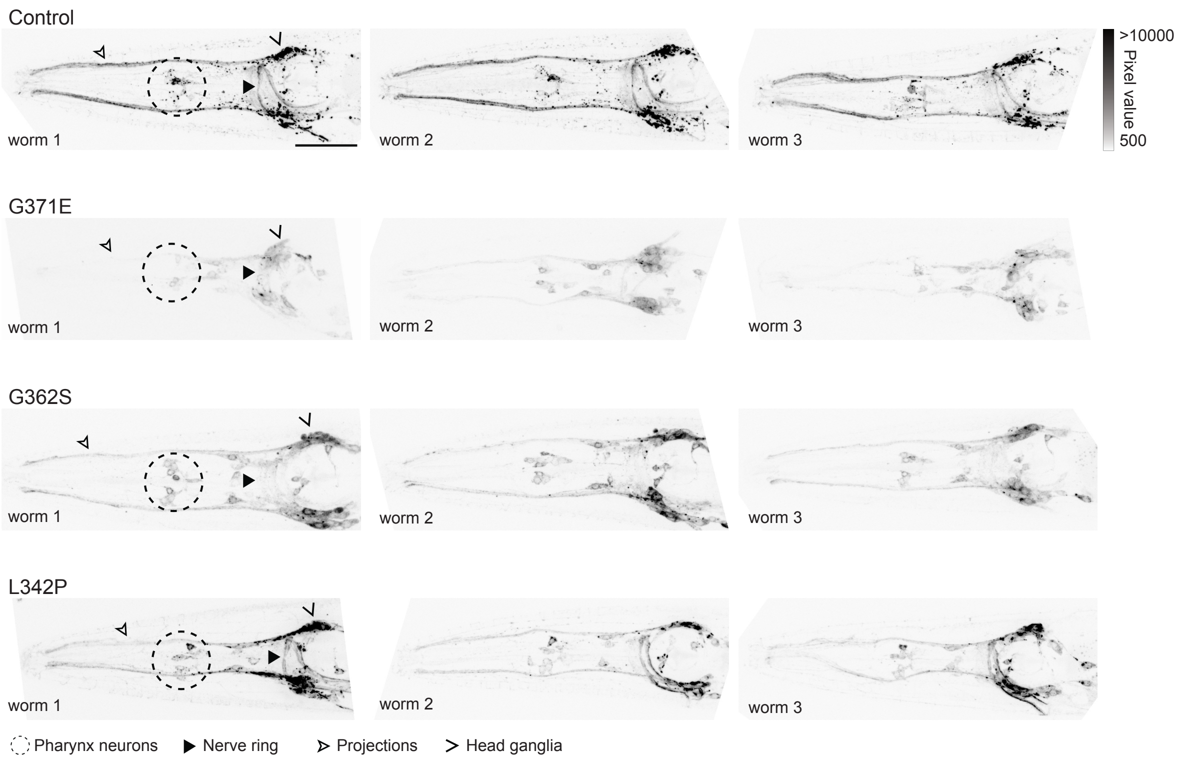

**Figure S4. Supplementary images for KCNL-1 fluorescence profiles in the head of control animals and of G371E, G362S, and L432P NEDMAB-hSK2 variants.**

Representative images of three different animals showing KCNL-1 localization in the head of control animals and G371E, G362S and L432P mutants. Fluorescence is visible in several neurons of the pharynx (dashed circle), the head ganglia (open arrowhead), projections to the tip of the nose (empty arrowhead), and the nerve ring (arrowhead). Dorso-ventral views. Scale bar, 20 µm.

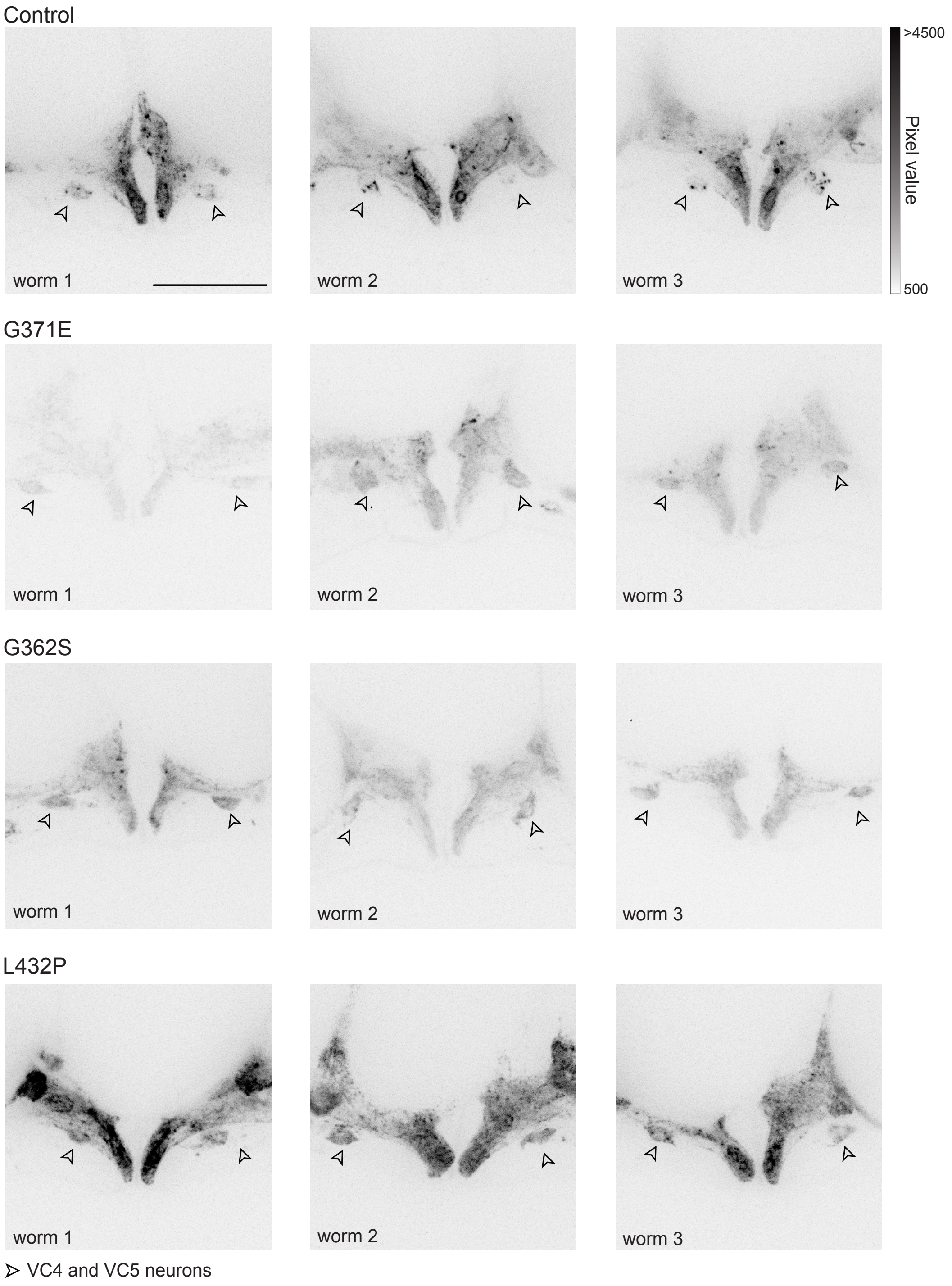

**Figure S5. Supplementary images for KCNL-1 fluorescence profiles in the egg-laying apparatus of control animals and of G371E, G362S, and L432P NEDMAB-hSK2 variants.**

Representative images of three different animals showing KCNL-1 localization in the egg-laying apparatus of control animals or mutants carrying the G371E, G362S, or L432P NEDMAB-hSK2 variants. Cell bodies of VC4 and VC5 motor neurons are indicated by empty arrowheads. Lateral views; anterior, left; ventral, down. Scale bar, 20 μm.

#### Supplementary table ST1

| Figure | Panel | Genotype | Strain | Allele | Description |
|---|---|---|---|---|---|
| Fig. 1 | C | Control (*kcnl-1* ::wScarlet) | JIP3105 | *bab315* | *kcnl-1* ::wrmScarlet; crRNA: oNB019 ; Repair template: from pSEM103 WITH oNB167/oNB168, PCR duplex for detention: oBR611-oNB165-oNB166 |
| | | *kcnl-1(0)* | JIP3131 | *bab315bln1118* | *kcnl-1* ::wrmScarlet indel 377_792del; crRNA: oMG054 ; Repair template: oMG174 ; PCR for detention: oMG175-oMG093-oMG88 |

| Figure | Panel | Patient variants | Strain | Allele | Description |
|---|---|---|---|---|---|
| Fig. 2 | B, C | V530L | JIP3106 | *bab315bab568* | *kcnl-1* ::wrmScarlet V530L ; crRNA: oMG029 ; Repair template: oMG083 ; PCR: oMG084/oMG085 |
| | | S436C | JIP3108 | *bab315bab567* | *kcnl-1* ::wrmScarlet S436C ; crRNA: MG030 ; Repair template: oMG086 ; PCR: oMG087/oMG088 |
| | | G350D | JIP3127 | *bab315bab618* | *kcnl-1* ::wrmScarlet G374D ; crRNA: oMG037 ; Repair template: oMG107 ; PCR: oMG110/oMG111 |
| Fig. 3 | B, C | G362S | JIP3121 | *bab315bab580* | *kcnl-1* ::wrmScarlet G511S ; crRNA: oMG034 ; Repair template: oMG105 ; PCR: oMG084/oMG085 |
| | | L432P | JIP3124 | *bab315bab595* | *kcnl-1* ::wrmScarlet F581P ; crRNA: oMG036 ; Repair template: oMG106 ; PCR: oMG109/oMG108 |
| | | L388V | JIP3123 | *bab315bab582* | *kcnl-1* ::wrmScarlet A537V ; crRNA: oMG035 ; Repair template: oMG103 ; PCR: oMG084/oMG085 |
| | | I288S | JIP3114 | *bab315bab616* | *kcnl-1* ::wrmScarlet I437S ; crRNA: MG030 ; Repair template: oMG090 ; PCR: oMG087/oMG088 |
| | | G371E | JIP3110 | *bab315bab613* | *kcnl-1* ::wrmScarlet G520E ; crRNA: MG032 ; Repair template: oMG089 ; PCR: oMG084/oMG085 |
| Fig. 4 | B, C | I359M | JIP3115 | *bab315 bab570* | *kcnl-1* ::wrmScarlet I508M ; crRNA: MG034 ; Repair template: oMG091 ; PCR: oMG084/oMG085 |
| Fig. 5 | B, C | V530L | JIP3106 | *bab315bab568* | *kcnl-1* ::wrmScarlet V530L ; crRNA: oMG029 ; Repair template: oMG083 ; PCR: oMG084/oMG085 |
| | | S436C | JIP3108 | *bab315bab567* | *kcnl-1* ::wrmScarlet S436C ; crRNA: MG030 ; Repair template: oMG086 ; PCR: oMG087/oMG088 |
| | | G350D | JIP3127 | *bab315bab618* | *kcnl-1* ::wrmScarlet G374D ; crRNA: oMG037 ; Repair template: oMG107 ; PCR: oMG110/oMG111 |
| | | I288S | JIP3114 | *bab315bab616* | *kcnl-1* ::wrmScarlet I437S ; crRNA: MG030 ; Repair template: oMG090 ; PCR: oMG087/oMG088 |
| | | I359M | JIP3115 | *bab315 bab570* | *kcnl-1* ::wrmScarlet I508M ; crRNA: MG034 ; Repair template: oMG091 ; PCR: oMG084/oMG085 |
| | | L388V | JIP3123 | *bab315bab582* | *kcnl-1* ::wrmScarlet A537V ; crRNA: oMG035 ; Repair template: oMG103 ; PCR: oMG084/oMG085 |
| | | G371E | JIP3110 | *bab315bab613* | *kcnl-1* ::wrmScarlet G520E ; crRNA: MG032 ; Repair template: oMG089 ; PCR: oMG084/oMG085 |
| | | G362S | JIP3121 | *bab315bab580* | *kcnl-1* ::wrmScarlet G511S ; crRNA: oMG034 ; Repair template: oMG105 ; PCR: oMG084/oMG085 |
| | | L432P | JIP3124 | *bab315bab595* | *kcnl-1* ::wrmScarlet F581P ; crRNA: oMG036 ; Repair template: oMG106 ; PCR: oMG109/oMG108 |
| Fig. 6 | A, B | G371E | JIP3110 | *bab315bab613* | *kcnl-1* ::wrmScarlet G520E ; crRNA: MG032 ; Repair template: oMG089 ; PCR: oMG084/oMG085 |
| | | G362S | JIP3121 | *bab315bab580* | *kcnl-1* ::wrmScarlet G511S ; crRNA: oMG034 ; Repair template: oMG105 ; PCR: oMG084/oMG085 |
| | | L432P | JIP3124 | *bab315bab595* | *kcnl-1* ::wrmScarlet F581P ; crRNA: oMG036 ; Repair template: oMG106 ; PCR: oMG109/oMG108 |
| Supp. Fig. 2 | A | V530L | JIP3107 | *bab315bab569* | *kcnl-1* ::wrmScarlet V530L ; crRNA: oMG029 ; Repair template: oMG083 ; PCR: oMG084/oMG085 |
| | | S436C | JIP3109 | *bab315bab566* | *kcnl-1* ::wrmScarlet S436C ; crRNA: MG030 ; Repair template: oMG086 ; PCR: oMG087/oMG088 |
| | B | G350D | JIP 3128 (line 2) | *bab315bab619* | *kcnl-1* ::wrmScarlet G374D ; crRNA: oMG037 ; Repair template: oMG107 ; PCR: oMG110/oMG111 |
| | | | JIP3129 (line 3) | *bab315bab620* | *kcnl-1* ::wrmScarlet G374D ; crRNA: oMG037 ; Repair template: oMG107 ; PCR: oMG110/oMG111 |
| | C | G362S | JIP3120 | *bab315bab579* | *kcnl-1* ::wrmScarlet G511S ; crRNA: oMG034 ; Repair template: oMG105 ; PCR: oMG084/oMG085 |
| | | L432P | JIP3126 | *bab315bab607* | *kcnl-1* ::wrmScarlet F581P ; crRNA: oMG036 ; Repair template: oMG106 ; PCR: oMG109/oMG108 |
| | | L388V | JIP3122 | *bab315bab581* | *kcnl-1* ::wrmScarlet A537V ; crRNA: oMG035 ; Repair template: oMG103 ; PCR: oMG084/oMG085 |
| | | I288S | JIP3113 | *bab315bab617* | *kcnl-1* ::wrmScarlet I437S ; crRNA: MG030 ; Repair template: oMG090 ; PCR: oMG087/oMG088 |
| | | G371E | JIP3111 | *bab315bab614* | *kcnl-1* ::wrmScarlet G520E ; crRNA: MG032 ; Repair template: oMG089 ; PCR: oMG084/oMG085 |
| | | I359M | JIP3116 | *bab315 bab571* | *kcnl-1* ::wrmScarlet I508M ; crRNA: MG034 ; Repair template: oMG091 ; PCR: oMG084/oMG085 |
| Supp. Fig. 3 | | G362S | JIP3121 | *bab315bab580* | *kcnl-1* ::wrmScarlet G511S ; crRNA: oMG034 ; Repair template: oMG105 ; PCR: oMG084/oMG085 |
| | | L432P | JIP3124 | *bab315bab595* | *kcnl-1* ::wrmScarlet F581P ; crRNA: oMG036 ; Repair template: oMG106 ; PCR: oMG109/oMG108 |
| | | L388V | JIP3123 | *bab315bab582* | *kcnl-1* ::wrmScarlet A537V ; crRNA: oMG035 ; Repair template: oMG103 ; PCR: oMG084/oMG085 |
| | | I288S | JIP3114 | *bab315bab616* | *kcnl-1* ::wrmScarlet I437S ; crRNA: MG030 ; Repair template: oMG090 ; PCR: oMG087/oMG088 |
| | | G371E | JIP3110 | *bab315bab613* | *kcnl-1* ::wrmScarlet G520E ; crRNA: MG032 ; Repair template: oMG089 ; PCR: oMG084/oMG085 |
| Supp. Fig. 4 | | G371E | JIP3110 | *bab315bab613* | *kcnl-1* ::wrmScarlet G520E ; crRNA: MG032 ; Repair template: oMG089 ; PCR: oMG084/oMG085 |
| | | G362S | JIP3121 | *bab315bab580* | *kcnl-1* ::wrmScarlet G511S ; crRNA: oMG034 ; Repair template: oMG105 ; PCR: oMG084/oMG085 |
| | | L432P | JIP3124 | *bab315bab595* | *kcnl-1* ::wrmScarlet F581P ; crRNA: oMG036 ; Repair template: oMG106 ; PCR: oMG109/oMG108 |
| Supp. Fig. 5 | | G371E | JIP3110 | *bab315bab613* | *kcnl-1* ::wrmScarlet G520E ; crRNA: MG032 ; Repair template: oMG089 ; PCR: oMG084/oMG085 |
| | | G362S | JIP3121 | *bab315bab580* | *kcnl-1* ::wrmScarlet G511S ; crRNA: oMG034 ; Repair template: oMG105 ; PCR: oMG084/oMG085 |
| | | L432P | JIP3124 | *bab315bab595* | *kcnl-1* ::wrmScarlet F581P ; crRNA: oMG036 ; Repair template: oMG106 ; PCR: oMG109/oMG108 |

### Supplementary table ST2

| crRNA name | Sequence (5'-3') |
| --- | --- |
| crRNA: oNB019 | GCTGATTTCAAATTTCGAAG |
| crRNA: MG029 | ATCTATTACTACTGGAATTG |
| crRNA: MG030 | AATCCGGTTCAGTGCTGCGA |
| crRNA MG032 | ATCGTCCCGAACACTTATTG |
| crRNA: MG034 | ACATTCATGTCTATCGGCTA |
| crRNA: oMG035 | ccagGGAGCCGGTGTCTCGT |
| crRNA: oMG036 | GCAGAGCTTTTTTGTACTTG |
| crRNA: oMG037 | AAAATAGCACTGATAGATAG |
| crRNA: oMG054 | TTCGAAGAaGTTGGAGCATC |

| Repair template name | Sequence (5'-3') |
| --- | --- |
| oMG083 | GCTCCctggaattgaaaaattttgcactttttttggcccaaactctagaaagcctacaAgAATTCCAGTAGTAATAGATAAACACCGTCCACAATAAGTGTTCGGGACGATATCTCCGTA |
| oMG086 | tgaaacttaaaaacggaaaaatctgaactttcgagttttcagGACGCCGCCACCCGGTgtATCGCAGCACTGAACCGGATTTCCATGGATTTTCGATTCGTGATCAAGACCATGATGGCT |
| oMG089 | ttttggcccaaactctagaaagcctacCACAATTCCAGTAGTAATAGATAAACAgCGTtCACAATAAGTGTTCGGGACGATATCTCCGTAGCCGATAGACATGAATGTGACCATTATGAA |
| oMG090 | aacttaaaaacggaaaaatctgaactttcgagttttcagGACGCCGCCACCCGGTCatcaGCAGCACTGAACCGGATTTCCATGGATTTTCGATTCGTGATCAAGACCATGATGGCTGAT |
| oMG031 | AGTAGTAATAGATAAACACCGTCCACAATAAGTGTTCGGGACGATATCTCCGTAGCCcATAGACATGAATGTGACCATTATGAACCACATCGAGTTGAGGTAGTAGTGCTCCACTTCCAG |
| oMG105 | TTCCAGTAGTAATAGATAAACACCGTCCACAATAAGTGTTCGGGACGATATCTgaGTAGCCGATAGACATGAATGTGACCATTATGAACCACATCGAGTTGAGGTAGTAGTGCTCCACTT |
| oMG103 | TGTTCACGTGCTTCTCGGCTCGTGAGAGCTCCAATTTTCGCGAAATAATTGCAATTAGAaCtGACGAGACACCGGCTCCctggaattgaaaaattttgcactttttttggcccaaactct |
| oMG106 | tccaaattctatggtctatttaaatttttctcaattttcctatttcagGccaATtCACAAGTACAAAAAAGCTCTGCACAAGGGTGACGACTTGCGACTGCGCCACCATCAACGCCGCTT |
| oMG107 | GATCAACATTTTAATAAATCGATCAGTACTGAACGCCACCCTCCAATCATCGGCatcACTATCTATCAGTGCTATTTTCACCTCGATTGCGTGATAATTGATGAGACAGCAGATAAGGAA |
| oMG174 | ttccaactgaagaaatgatcagcgcacaattttacattcaacatcttgctttgttttcagGGAGCATCGGGAGCCTCAGGAGCATCGATGGTCAGCAAGGGAGAGGCAGTTATCAAGGAG |

| Oligo name (PCR) | Sequence (5'-3') |
| --- | --- |
| oMG084 | GCATGTCGTGGATGTTCACC |
| oMG085 | TGCGGCATTCTTCCGTTGAT |
| oMG087 | GACGGGCTCATGAAAATGCC |
| oMG088 | ggcttcaaggggacttacCG |
| oMG108 | GTACGGTGCATCTCCCAGAG |
| oMG109 | cctatacgctctctcaccgc |
| oMG110 | CGGTACTCGGCCTCCTATTG |
| oMG111 | ggcggcgatatagtgtggaa |
| oMG175 | ctctggcgaccttctcttgg |
| oMG088 | ggcttcaaggggacttacCG |
| oMG093 | CCTCTCCCCACAATTCATGTACG |
| oBR611 | AGGAGAATGGGAGTGGTCCT |
| oNB165 | CAGGTCACCACCACGACTAC |
| oNB166 | aatgctgcctctctgtacgc |

#### Supplementary table ST3

| hSK3 varian | KCNL-1 variant |
|---|---|
| Ser436Cys | Ser436Cys |
| Gly350Asp | Gly347Asp |

| hSK2 varian | KCNL-1 variant |
|---|---|
| Gly371Glu | Gly520Glu |
| Ile288Ser | Ile437Ser |
| Ile359Met | Ile508Met |
| Gly362Ser | Gly511Ser |
| Leu388Val | Ala537Val |
| Leu432Pro | Phe581Pro |

#### Supplementary table ST4

| Figure | Panel | Assay | Variants | Statistical test | P value |
|---|---|---|---|---|---|
| Fig. 1 | C | Embryos in utero | *kcnl-1(0)* | Mann Whitney test | <0.0001 |
| | | Embryonic stage (%) | | Fisher's Exact Test and Bonferroni correction | <20 cells p = 0.0178<br>20 cells to gastrula p = 0.0178 |
| Fig. 2 | C | Embryos in utero | V530L | Mann Whitney test | <0.0001 |
| | | | S436C | Mann Whitney test | <0.0001 |
| | | | G350D | Mann Whitney test | 0.0012 |
| | | Embryonic stage (%) | V530L | Fisher's Exact Test and Bonferroni correction | <20 cells p = 0.000;<br>20 cells to gastrula p = 0.5018;<br>post-gastrula p = 0.000 |
| | | | S436C | Fisher's Exact Test and Bonferroni correction | <20 cells p = 0.000;<br>20 cells to gastrula p = 1.000;<br>post-gastrula p = 0.000 |
| | | | G350D | Fisher's Exact Test and Bonferroni correction | <20 cells p = 0.000;<br>20 cells to gastrula p = p = 0.000; |
| | | Offspring (72 h) | V530L | Mann Whitney test | <0.0001 |
| | | | S436C | Mann Whitney test | 0.0172 |
| | | | G350D | Mann Whitney test | 0.0559 |
| Fig. 3 | C | Embryos in utero | G362S | Kruskal-Wallis test | <0.0001 |
| | | | L432P | Kruskal-Wallis test | <0.0001 |
| | | | L388V | Kruskal-Wallis test | P value Control vs. L388V 0,0906; P value L388V vs. *kcnl-1(0)* 0,0672 |
| | | | I288S | Kruskal-Wallis test | <0.0001 |
| | | | G371E | Kruskal-Wallis test | <0.0001 |
| | | Embryonic stage (%) | G362S | Fisher's Exact Test and Bonferroni correction | **Control vs G362S**<br>• <20 cells p=0.0391<br>• 20 cells to gastrula p=0.0391<br>**Control vs *kcnl-1(0)***<br>• <20 cells p=0.0817<br>• 20 cells to gastrula p=0.0817<br>***kcnl-1(0)* vs G362S**<br>• <20 cells p=1<br>• 20 cells to gastrula p=1 |
| | | | L432P | Fisher's Exact Test and Bonferroni correction | **Control vs L432P**<br>• <20 cells p=0.0053<br>•20 cells to gastrula p=0.0053<br>**Control vs *kcnl-1(0)***<br>• <20 cells p=0.0004<br>• 20 cells to gastrula p=0.0004<br>***kcnl-1(0)* vs L432P**<br>• <20 cells p=0.8139<br>• 20 cells to gastrula p=0.8139 |
| | | | L388V | Fisher's Exact Test and Bonferroni correction | **Control vs L388V**<br>• <20 cells p=0.0019<br>•20 cells to gastrula p=0.0019<br>**Control vs *kcnl-1(0)***<br>• <20 cells p=0.0038<br>• 20 cells to gastrula p=0.0038<br>***kcnl-1(0)* vs L388V**<br>• <20 cells p=1<br>• 20 cells to gastrula p=1 |
| | | | I288S | Fisher's Exact Test and Bonferroni correction | **Control vs I288S**<br>• <20 cells p=0.0003<br>•20 cells to gastrula p=0.0003<br>**Control vs *kcnl-1(0)***<br>• <20 cells p=0.0040<br>• 20 cells to gastrula p=0.0040<br>***kcnl-1(0)* vs I288S**<br>• <20 cells p=1<br>• 20 cells to gastrula p=1 |
| | | | G371E | Fisher's Exact Test and Bonferroni correction | **Control vs G371E**<br>• <20 cells p=0.0009<br>•20 cells to gastrula p=0.0009<br>**Control vs *kcnl-1(0)***<br>• <20 cells p=0.0040<br>• 20 cells to gastrula p=0.0040<br>***kcnl-1(0)* vs G371E**<br>• <20 cells p=1<br>• 20 cells to gastrula p=1 |
| Fig. 4 | C | Embryos in utero | I359M | Ordinary One-Way ANOVA | <0.0001 |
| | | Embryonic stage (%) | | Fisher's Exact Test and Bonferroni correction | **Control vs I359M**<br>• <20 cells p=0.0000<br>• 20 cells to gastrula p=0.9533<br>• post gastrula p=0.000<br>**Control vs *kcnl-1(0)***<br>• <20 cells p=0.0000<br>• 20 cells to gastrula p=0.0000<br>• post gastrula p=1<br>***kcnl-1(0)* vs I359M**<br>• <20 cells p=0.000<br>• 20 cells to gastrula p=0.000<br>• post gastrula p=0.000 |
| | | Offspring (72 h) | | Kruskal-Wallis test | <0.0001 |
| Supp. Fig. 2 | A | Embryos in utero | V530L | Mann Whitney test | <0.0001 |
| | | | S436C | Mann Whitney test | <0.0001 |
| | B | Embryos in utero | G350D line 2 JIP3128) | Mann Whitney test | 0.1930 |
| | | | G350D line 3 (JIP3129) | Mann Whitney test | 0.3307 |
| | | Embryonic stage (%) | G350D line 2 JIP3128) | Fisher's Exact Test and Bonferroni correction | <20 cells p = 0.6650<br>20 cells to gastrula p = 0.6650 |
| | | | G350D line 3 (JIP3129) | Fisher's Exact Test and Bonferroni correction | <20 cells p = 0.000<br>20 cells to gastrula p = 0.0000 |
| | C | Embryos in utero | G362S | Kruskal-Wallis test | <0.0001 |
| | | | L432P | Kruskal-Wallis test | 0.0007 |
| | | | L388V | Kruskal-Wallis test | <0.0001 |
| | | | I288S | Kruskal-Wallis test | <0.0001 |
| | | | G371E | Ordinary One-Way ANOVA | <0.0001 |
| | | | I359M | Ordinary One-Way ANOVA | <0.0001 |
| Supp. Fig.3 | | Brood size | G362S | Kruskal-Wallis test | 0.8981 |
| | | | L432P | Kruskal-Wallis test | 0.9251 |
| | | | L388V | Ordinary One-Way ANOVA | 0.4123 |
| | | | I288S | Ordinary One-Way ANOVA | 0.0307 |
| | | | G371E | Ordinary One-Way ANOVA | 0.1368 |

### Appendix 1

#### Figure 1C

##### Embryos in utero - Control vs *kcnl-1(0)*

First repetition | Second repetition

| Control (JIP3105) | *kcnl-1(0)* JIP3131 | Control (JIP3105) | G350D (JIP3127) |
|---|---|---|---|
| 13 | 9 | 11 | 9 |
| 13 | 14 | 10 | 8 |
| 16 | 12 | 16 | 14 |
| 13 | 11 | 17 | 10 |
| 10 | 8 | 12 | 12 |
| 11 | 10 | 12 | 8 |
| 12 | 9 | 18 | 7 |
| 13 | 6 | 13 | 11 |
| 10 | 9 | 9 | 7 |
| 18 | 12 | 15 | 6 |
| 10 | 13 | 13 | 13 |
| 12 | 8 | 11 | 9 |
| 16 | 12 | 12 | 8 |
| 14 | 8 | 9 | 9 |
| 16 | 9 | 18 | 9 |
| 13 | 11 | 11 | 12 |
| 15 | 10 | 16 | 10 |
| 16 | 9 | 8 | 8 |
| 15 | 8 | 14 | 12 |
| 9 | 7 | 16 | 10 |
| 13 | 10 | 17 | 9 |
| 14 | 11 | 14 | 10 |
| 16 | 8 | 14 | 7 |
| 12 | 9 | 12 | 9 |
| 12 | 13 | 18 | 11 |
| 16 | 6 | 16 | 9 |
| 13 | 10 | 13 | 5 |
| | 13 | 11 | 12 |
| | 7 | 10 | 10 |
| | 8 | | |

Mann Whitney test
P value <0.0001

##### Embryonic stage (%) - Control vs *kcnl-1(0)*

| | <20 cells | 20 cells to gastrula |
|---|---|---|
| **Control (JIP3105)** | 29 | 71 |
| **kcnl-1(0) (JIP3131)** | 48 | 53 |

Fisher's Exact + Bonferroni correction

| **Category** | **P value** | **Correction** |
|---|---|---|
| <20 cells | 0.0089 | 0.0178 |
| 20 cells to gastrula | 0.0089 | 0.0178 |

**Figure 2C**

#### Embryos in utero - V530L and ZLS variants

| First repetition | | Second repetition | | First repetition | | Second repetition | |
|---|---|---|---|---|---|---|---|
| Control (JIP3105) | V530L (JIP3106) | Control (JIP3105) | V530L (JIP3106) | Control (JIP3105) | S436C (JIP3108) | Control (JIP3105) | S436C (JIP3108) |
| 12 | 32 | 24 | 50 | 19 | 37 | 12 | 47 |
| 15 | 37 | 23 | 45 | 18 | 40 | 21 | 43 |
| 19 | 31 | 21 | 43 | 15 | 40 | 16 | 57 |
| 14 | 23 | 30 | 23 | 13 | 36 | 21 | 45 |
| 17 | 30 | 19 | 29 | 16 | 38 | 12 | 46 |
| 19 | 35 | 15 | 34 | 28 | 40 | 23 | 42 |
| 20 | 26 | 28 | 44 | 19 | 38 | 20 | 46 |
| 19 | 28 | 21 | 46 | 14 | 36 | 21 | 42 |
| 20 | 25 | 18 | 36 | 18 | 35 | 28 | 33 |
| 15 | 20 | 22 | 36 | 14 | 32 | 24 | 32 |
| 15 | 35 | 26 | 57 | 14 | 31 | 20 | 45 |
| 22 | 32 | 21 | 31 | 14 | 50 | 20 | 59 |
| 16 | 44 | 19 | 38 | 19 | 42 | 17 | 48 |
| 22 | 22 | 16 | 30 | 18 | 39 | 12 | 60 |
| 19 | 30 | 21 | 36 | 17 | 39 | 22 | 56 |
| 16 | 18 | 21 | 29 | 25 | 42 | 21 | 36 |
| 16 | 18 | 19 | 31 | 13 | 41 | 32 | 38 |
| 20 | 26 | 22 | 41 | 16 | 36 | 18 | 58 |
| 10 | 34 | 23 | 37 | 15 | 39 | 35 | 38 |
| 13 | 20 | 16 | 41 | 16 | 39 | 26 | 47 |
| 17 | 34 | 21 | 52 | 12 | 39 | 21 | 56 |
| 17 | 20 | 21 | 39 | 16 | 39 | 25 | 42 |
| 22 | 41 | 22 | 37 | 20 | 25 | 20 | 43 |
| 13 | 38 | 23 | 26 | 15 | 37 | 24 | 59 |
| 22 | 27 | 22 | 31 | 18 | 18 | 18 | 47 |
| 14 | 30 | 24 | 36 | 21 | 39 | 27 | 51 |
| 13 | 26 | 14 | 52 | 19 | 42 | 20 | 49 |
| 19 | 40 | 21 | 40 | 22 | 19 | 13 | 44 |
| 11 | 26 | 20 | 49 | 15 | 40 | 18 | 28 |
| | 22 | 25 | 37 | 16 | 31 | 15 | 44 |
| Mann Whitney test P value <0.0001 | | | | Mann Whitney test P value <0.0001 | | | |

**Figure 2C**

#### Embryos in utero - V530L and ZLS variants

| First repetition | | Second repetition | |
|---|---|---|---|
| Control (JIP3105) | G350D (JIP3127) | Control (JIP3105) | G350D (JIP3127) |
| 18 | 18 | 21 | 30 |
| 20 | 20 | 16 | 20 |
| 19 | 22 | 20 | 26 |
| 16 | 21 | 18 | 23 |
| 18 | 21 | 30 | 32 |
| 13 | 17 | 20 | 18 |
| 25 | 34 | 27 | 29 |
| 19 | 16 | 38 | 24 |
| 15 | 19 | 16 | 30 |
| 17 | 20 | 16 | 30 |
| 17 | 19 | 19 | 17 |
| 14 | 26 | 25 | 31 |
| 15 | 20 | 12 | 24 |
| 22 | 19 | 27 | 26 |
| 20 | 16 | 20 | 21 |
| 15 | 20 | 21 | 29 |
| 16 | 20 | 24 | 22 |
| 18 | 21 | 32 | 29 |
| 16 | 19 | 13 | 22 |
| 16 | 35 | 19 | 14 |
| 18 | 20 | 14 | 21 |
| 20 | 16 | 28 | 25 |
| 16 | 20 | 20 | 17 |
| 19 | 19 | 16 | 10 |
| 22 | 14 | 18 | 26 |
| 27 | 23 | 20 | 14 |
| 21 | 22 | 21 | 19 |
| 19 | 21 | 21 | 22 |
| 21 | 23 | 15 | 27 |
| 15 | 19 | 17 | 33 |

Mann Whitney test
P value 0.0012

**Figure 2C**

#### Embryonic stage (%) - V530L and ZLS variants

| | <20 cells | 20 cells to gastrula | post gastrula |
|---|---|---|---|
| **Control (JIP3105)** | 34 | 68 | 0 |
| **V530L (JIP3106)** | 0 | 78 | 25 |

**Fisher's Exact + Bonferroni correction**

| Category | P value | Correction |
|---|---|---|
| <20 cells | 0.0000 | 0.0000 |
| 20 cells to gastrula | 0.1673 | 0.5018 |
| post gastrula | 0.0000 | 0.0000 |

| | <20 cells | 20 cells to gas | post gastrula |
|---|---|---|---|
| **Control (JIP3105)** | 38 | 62 | 0 |
| **S436C (JIP3108)** | 0 | 63 | 40 |

**Fisher's Exact + Bonferroni correction**

| Category | P value | Correction |
|---|---|---|
| <20 cells | 0.0000 | 0.0000 |
| 20 cells to gastrula | 1.0000 | 1.0000 |
| post gastrula | 0.0000 | 0.0000 |

| | <20 cells | 20 cells to gas | post gastrula |
|---|---|---|---|
| **Control (JIP3105)** | 50 | 68 | 0 |
| **G350D (JIP3127)** | 10 | 97 | 0 |

**Fisher's Exact + Bonferroni**

| Category | P value | Correction |
|---|---|---|
| <20 cells | 0.0000 | 0.0000 |
| 20 cells to gastrula | 0.0000 | 0.0000 |

**Figure 2C**

#### Brood size - V530L and ZLS variants

| Control (JIP3105) | V530L (JIP3106) |
|---|---|
| 276 | 172 |
| | 216 |
| 236 | 222 |
| 260 | 187 |
| 267 | 102 |
| 278 | 38 |
| 283 | 182 |
| 310 | 204 |
| 259 | 173 |
| 250 | 171 |

Mann Whitney test
P value <0.0001

| Control (JIP3105) | S436C (JIP3108) |
|---|---|
| 350 | 254 |
| | 231 |
| 303 | 48 |
| 343 | 199 |
| 195 | 244 |
| 145 | 236 |
| 338 | 217 |
| 326 | 72 |
| 269 | 211 |
| 352 | 202 |

Mann Whitney test
P value = 0.0172

| Control (JIP3105) | G350D (JIP3127) |
|---|---|
| 316 | 351 |
| 290 | 293 |
| 285 | 252 |
| 340 | 268 |
| 331 | 288 |
| 345 | 241 |
| 324 | 331 |
| 323 | 288 |
| 340 | 322 |
| | 275 |

Mann Whitney test
P value = 0.0559

**Figure 3C**

#### Embryos in utero - NEDMAB variants

| First repetition | | | Second repetition | | |
|---|---|---|---|---|---|
| Control (JIP3105) | G362S (JIP3121) | *kcnl-1(0)* (JIP3131) | Control (JIP3105) | G362S (JIP3121) | *kcnl-1(0)* (JIP3131) |
| 21 | 17 | 11 | 27 | 21 | 18 |
| 17 | 14 | 15 | 30 | 11 | 15 |
| 23 | 13 | 17 | 13 | 14 | 27 |
| 16 | 12 | 13 | 24 | 5 | 16 |
| 23 | 16 | 17 | 18 | 18 | 17 |
| 17 | 10 | 15 | 27 | 20 | 19 |
| 21 | 20 | 12 | 26 | 17 | 17 |
| 21 | 17 | 15 | 24 | 16 | 9 |
| 21 | 5 | 15 | 18 | 13 | 12 |
| 23 | 7 | 10 | 25 | 18 | 13 |
| 27 | 12 | 10 | 24 | 11 | 11 |
| 21 | 14 | 11 | 27 | 13 | 17 |
| 15 | 11 | 14 | 19 | 13 | 9 |
| 15 | 11 | 11 | 12 | 8 | 17 |
| 22 | 11 | 17 | 14 | 20 | 14 |
| 23 | 16 | 13 | 15 | 14 | 11 |
| 13 | 8 | 13 | 28 | 13 | 28 |
| 28 | 8 | 12 | 25 | 19 | 15 |
| 21 | 7 | 9 | 12 | 13 | 13 |
| 22 | 8 | 17 | 7 | 15 | 14 |
| 19 | 16 | 12 | 23 | 14 | 11 |
| 16 | 11 | 9 | 29 | 10 | 19 |
| 20 | 13 | 11 | 20 | 17 | 16 |
| 20 | 6 | 6 | 18 | 19 | 15 |
| 17 | 17 | 7 | 25 | 16 | 16 |
| 16 | 17 | 11 | 22 | 10 | 29 |
| 19 | 26 | 10 | 18 | 10 | 12 |
| 16 | 8 | 15 | 16 | 17 | 14 |
| 27 | 9 | 9 | 19 | 17 | 18 |
| 24 | 11 | 18 | 21 | 20 | 11 |

Kruskal-Wallis test P value <0.0001

**Figure 3C**

#### Embryos in utero - NEDMAB variants

First repetition / Second repetition

| Control (JIP3105) | L432P (JIP3124) | *kcnl-1(0)* (JIP3131) | Control (JIP3105) | L432P (JIP3124) | *kcnl-1(0)* (JIP3131) |
|---|---|---|---|---|---|
| 17 | 8 | 12 | 16 | 19 | 14 |
| 11 | 18 | 8 | 30 | 13 | 15 |
| 22 | 17 | 12 | 17 | 19 | 16 |
| 16 | 11 | 17 | 16 | 11 | 12 |
| 19 | 11 | 10 | 11 | 16 | 17 |
| 27 | 13 | 10 | 21 | 16 | 15 |
| 16 | 16 | 13 | 17 | 20 | 11 |
| 15 | 17 | 9 | 25 | 13 | 14 |
| 17 | 19 | 12 | 18 | 10 | 17 |
| 15 | 14 | 12 | 16 | 12 | 19 |
| 26 | 16 | 12 | 24 | 8 | 9 |
| 18 | 14 | 10 | 17 | 12 | 12 |
| 16 | 13 | 23 | 22 | 14 | 11 |
| 17 | 10 | 10 | 23 | 12 | 18 |
| 23 | 22 | 25 | 17 | 14 | 15 |
| 14 | 12 | 14 | 22 | 11 | 18 |
| 9 | 13 | 16 | 11 | 16 | 14 |
| 17 | 14 | 27 | 18 | 12 | 13 |
| 24 | 13 | 14 | 23 | 12 | 13 |
| 19 | 8 | 10 | 17 | 11 | 10 |
| 19 | 12 | | 19 | 13 | 13 |
| 16 | 16 | | 21 | 13 | 13 |
| 21 | 12 | | 21 | 15 | 15 |
| 12 | 13 | | 20 | 15 | 6 |
| 14 | 17 | | 20 | 15 | 19 |
| 15 | 21 | | 17 | 13 | 15 |
| 16 | 17 | | 15 | 15 | 8 |
| 17 | 13 | | 20 | 9 | 27 |
| 17 | 15 | | 22 | 12 | 10 |
| 12 | 22 | | 16 | 15 | 11 |

Kruskal-Wallis test P value <0.0001

**Figure 3C**

#### Embryos in utero - NEDMAB variants

First repetition | Second repetition

| Control (JIP3105) | L388V (JIP3123) | *kcnl-1(0)* (JIP3131) | Control (JIP3105) | L388V (JIP3123) | *kcnl-1(0)* (JIP3131) |
|---|---|---|---|---|---|
| 20 | 22 | 16 | 12 | 13 | 11 |
| 12 | 6 | 22 | 14 | 10 | 8 |
| 20 | 18 | 18 | 16 | 12 | 6 |
| 16 | 20 | 18 | 18 | 17 | 9 |
| 19 | 15 | 15 | 18 | 12 | 11 |
| 15 | 19 | 23 | 12 | 10 | 8 |
| 24 | 17 | 14 | 11 | 10 | 9 |
| 16 | 16 | 8 | 11 | 12 | 10 |
| 25 | 18 | 8 | 14 | 10 | 7 |
| 18 | 20 | 17 | 12 | 11 | 13 |
| 11 | 18 | 9 | 16 | 9 | 17 |
| 13 | 18 | 11 | 14 | 16 | 8 |
| 22 | 15 | 13 | 12 | 9 | 11 |
| 17 | 16 | 11 | 13 | 7 | 10 |
| 13 | 18 | 1 | 16 | 8 | 14 |
| 23 | 16 | 21 | 10 | 11 | 5 |
| 18 | 15 | 17 | 14 | 12 | 7 |
| 10 | 14 | 11 | 12 | 11 | 15 |
| 22 | 18 | 23 | 12 | 11 | 8 |
| 21 | 28 | 13 | 12 | 8 | 7 |
| 18 | 6 | 18 | 13 | 14 | 9 |
| 16 | 14 | 14 | 12 | 9 | 6 |
| 14 | 18 | 14 | 10 | 11 | 8 |
| 22 | 13 | 11 | 10 | 14 | 4 |
| 28 | 16 | 14 | 10 | 12 | 9 |
| 12 | 16 | 11 | 13 | 10 | 11 |
| 19 | 15 | 5 | 15 | 10 | 11 |
| 16 | 18 | 15 | 13 | 12 | 5 |
| 18 | 15 | 12 | 16 | 10 | 15 |
| 14 | 13 | 22 | | 10 | 7 |
| | | | | 8 | |

Kruskal-Wallis test P value Control vs. L388V 0,0906;
P value L388V vs. *kcnl-1(0)* 0,0672

**Figure 3C**

#### Embryos in utero - NEDMAB variants

First repetition / Second repetition

| Control (JIP3105) | I288S (JIP3114) | *kcnl-1(0)* (JIP3131) | Control (JIP3105) | I288S (JIP3114) | *kcnl-1(0)* (JIP3131) |
|---|---|---|---|---|---|
| 16 | 9 | 13 | 22 | 23 | 14 |
| 13 | 9 | 12 | 28 | 19 | 13 |
| 16 | 8 | 15 | 30 | 20 | 18 |
| 14 | 14 | 13 | 18 | 14 | 14 |
| 18 | 7 | 10 | 26 | 20 | 26 |
| 21 | 10 | 7 | 28 | 19 | 15 |
| 23 | 13 | 15 | 25 | 15 | 17 |
| 14 | 5 | 13 | 16 | 34 | 24 |
| 20 | 14 | 10 | 23 | 22 | 13 |
| 24 | 13 | 17 | 22 | 22 | 17 |
| 20 | 9 | 11 | 29 | 16 | 16 |
| 14 | 11 | 8 | 27 | 11 | 18 |
| 17 | 9 | 8 | 21 | 25 | 21 |
| 16 | 10 | 9 | 27 | 21 | 16 |
| 12 | 10 | 7 | 20 | 13 | 16 |
| 13 | 9 | 15 | 24 | 17 | 25 |
| 16 | 8 | 5 | 22 | 25 | 23 |
| 23 | 5 | 8 | 27 | 22 | 22 |
| 18 | 9 | 15 | 22 | 15 | 23 |
| 15 | 7 | 9 | 27 | 23 | 19 |
| 18 | 12 | 5 | 24 | 20 | 29 |
| 16 | 9 | 18 | 26 | 12 | 14 |
| 10 | 12 | 9 | 31 | 21 | 8 |
| 22 | 10 | 10 | 30 | 12 | 17 |
| 21 | 13 | 12 | 21 | 17 | 12 |
| 23 | 9 | 10 | 27 | 15 | 22 |
| 17 | 11 | 9 | 29 | 11 | 27 |
| 16 | 13 | 13 | 23 | 15 | 24 |
| 17 | 10 | 8 | 24 | 17 | 20 |
| 19 | 11 | 6 | 27 | 19 | 17 |

Kruskal-Wallis test P value <0.0001

#### Figure 3C

#### Embryos in utero - NEDMAB variants

| First repetition | | | Second repetition | | |
|---|---|---|---|---|---|
| Control (JIP3105) | G371E (JIP3110) | *kcnl-1(0)* (JIP3131) | Control (JIP3105) | G371E (JIP3110) | *kcnl-1(0)* (JIP3131) |
| 19 | 21 | 18 | 17 | 13 | 10 |
| 14 | 18 | 17 | 16 | 11 | 12 |
| 21 | 16 | 20 | 17 | 12 | 10 |
| 13 | 19 | 16 | 11 | 12 | 9 |
| 19 | 12 | 17 | 16 | 7 | 12 |
| 13 | 16 | 17 | 10 | 9 | 5 |
| 13 | 13 | 16 | 19 | 13 | 12 |
| 24 | 19 | 14 | 15 | 13 | 7 |
| 19 | 18 | 17 | 15 | 7 | 9 |
| 18 | 16 | 20 | 11 | 10 | 5 |
| 18 | 14 | 9 | 18 | 10 | 13 |
| 17 | 15 | 16 | 23 | 14 | 10 |
| 16 | 16 | 19 | 13 | 7 | 6 |
| 23 | 7 | 12 | 11 | 15 | 11 |
| 22 | 13 | 18 | 12 | 11 | 8 |
| 21 | 11 | 10 | 22 | 10 | 6 |
| 12 | 11 | 11 | 13 | 13 | 7 |
| 24 | 17 | 8 | 13 | 17 | 5 |
| 11 | 10 | 12 | 16 | 9 | 6 |
| 16 | 9 | 14 | 15 | 8 | 14 |
| 21 | 9 | 7 | 15 | 11 | 10 |
| 19 | 8 | 18 | 16 | 10 | 11 |
| 22 | 11 | 14 | 10 | 8 | 6 |
| 18 | 12 | 22 | 14 | 11 | 12 |
| 17 | 15 | 20 | 12 | 7 | 10 |
| 16 | 17 | 12 | 13 | 7 | 77 |
| 19 | 13 | 16 | 19 | 13 | 12 |
| 18 | 16 | 19 | 12 | 8 | 9 |
| 21 | 14 | 14 | 12 | 9 | |
| 23 | 14 | 18 | 15 | 8 | |

Kruskal-Wallis test P value <0.0001

#### Figure 3C

##### Embryonic stage (%) - NEDMAB variants

| | <20 cells | 20 cells to gastrula |
|---|---|---|
| **Control (JIP3105)** | 41 | 67 |
| **G362S (JIP3121)** | 58 | 48 |
| **kcnl-1(0) (JIP3131)** | 57 | 52 |

| Fisher Exact + Bonferroni correction | | | **Control vs G362S** | **Control vs *kcnl-1 (0)*** | ***kcnl-1 (0)* vs G362S** |
|---|---|---|---|---|---|
| | **< 20 cells** | P value | 0.0196 | 0.0409 | 0.7849 |
| | | Correction | 0.0391 | 0.0817 | 1.0000 |
| | **20 cells to gastrula** | P value | 0.0196 | 0.0409 | 0.7849 |
| | | Correction | 0.0391 | 0.0817 | 1.0000 |

| | <20 cells | 20 cells to gastrula |
|---|---|---|
| **Control (JIP3105)** | 28 | 74 |
| **L432P (JIP3124)** | 51 | 55 |
| **kcnl-1(0) (JIP3131)** | 54 | 46 |

| Fisher Exact + Bonferroni correction | | | **Control vs L432P** | **Control vs *kcnl-1 (0)*** | ***kcnl-1 (0)* vs L432P** |
|---|---|---|---|---|---|
| | **< 20 cells** | P value | 0.0027 | 0.0002 | 0.4069 |
| | | Correction | 0.0053 | 0.0004 | 0.8139 |
| | **20 cells to gastrula** | P value | 0.0027 | 0.0002 | 0.4069 |
| | | Correction | 0.0053 | 0.0004 | 0.8139 |

| | <20 cells | 20 cells to gastrula |
|---|---|---|
| **Control (JIP3105)** | 35 | 66 |
| **L388V (JIP3123)** | 63 | 47 |
| **kcnl-1(0) (JIP3131)** | 58 | 44 |

| Fisher Exact + Bonferroni correction | | | **Control vs L388V** | **Control vs *kcnl-1 (0)*** | ***kcnl-1 (0)* vs L388V** |
|---|---|---|---|---|---|
| | **< 20 cells** | P value | 0.0014 | 0.0019 | 1.0000 |
| | | Correction | 0.0029 | 0.0038 | 1.0000 |
| | **20 cells to gastrula** | P value | 0.0014 | 0.0019 | 1.0000 |
| | | Correction | 0.0029 | 0.0038 | 1.0000 |

#### Figure 3C

##### Embryonic stage (%) - NEDMAB variants

| | <20 cells | 20 cells to gastrula |
|---|---|---|
| **Control (JIP3105)** | 33 | 73 |
| **I288S (JIP3114)** | 64 | 48 |
| **kcnl-1(0) (JIP3131)** | 53 | 48 |

| Fisher Exact + Bonferroni correction | | | **Control vs I288S** | **Control vs *kcnl-1 (0)*** | ***kcnl-1 (0)* vs I288S** |
|---|---|---|---|---|---|
| | **< 20 cells** | P value | 0.0001 | 0.0020 | 0.5814 |
| | | Correction | 0.0003 | 0.0040 | 1.0000 |
| | **20 cells to gastrula** | P value | 0.0001 | 0.0020 | 0.5814 |
| | | Correction | 0.0001 | 0.0040 | 1.0000 |

| | <20 cells | 20 cells to gastrula |
|---|---|---|
| **Control (JIP3105)** | 34 | 69 |
| **G371E (JIP3110)** | 59 | 43 |
| **kcnl-1(0) (JIP3131)** | 65 | 56 |

| Fisher Exact + Bonferroni correction | | | **Control vs G371E** | **Control vs *kcnl-1 (0)*** | ***kcnl-1 (0)* vs G371E** |
|---|---|---|---|---|---|
| | **< 20 cells** | P value | 0.0004 | 0.0020 | 0.5892 |
| | | Correction | 0.0009 | 0.0040 | 1.0000 |
| | **20 cells to gastrula** | P value | 0.0004 | 0.0020 | 0.5892 |
| | | Correction | 0.0009 | 0.0040 | 1.0000 |

#### Figure 4C

##### Embryos in utero - I359M

| First repetition | | | Second repetition | | |
|---|---|---|---|---|---|
| **Control (JIP3105)** | **I359M (JIP3115)** | ***kcnl-1(0)* (JIP3131)** | **Control (JIP3105)** | **I359M (JIP3115)** | ***kcnl-1(0)* (JIP3131)** |
| 14 | 27 | 9 | 28 | 42 | 16 |
| 15 | 29 | 10 | 20 | 40 | 10 |
| 13 | 32 | 8 | 19 | 41 | 17 |
| 17 | 27 | 12 | 22 | 36 | 20 |
| 12 | 23 | 16 | 22 | 41 | 19 |
| 24 | 33 | 15 | 17 | 40 | 17 |
| 11 | 29 | 12 | 23 | 35 | 16 |
| 25 | 20 | 15 | 22 | 36 | 14 |
| 18 | 23 | 12 | 25 | 45 | 16 |
| 22 | 26 | 16 | 20 | 37 | 22 |
| 16 | 19 | 15 | 13 | 38 | 16 |
| 20 | 21 | 8 | 17 | 43 | 13 |
| 20 | 33 | 13 | 22 | 47 | 15 |
| 19 | 22 | 9 | 9 | 35 | 23 |
| 14 | 27 | 19 | 19 | 40 | 15 |
| 14 | 23 | 13 | 27 | 43 | 15 |
| 20 | 27 | 8 | 17 | 48 | 19 |
| 12 | 20 | 10 | 17 | 39 | 19 |
| 14 | 29 | 9 | 25 | 33 | 17 |
| 17 | 26 | 10 | 20 | 37 | 10 |
| 17 | 26 | 22 | 23 | 43 | 11 |
| 12 | 14 | 8 | 25 | 30 | 17 |
| 12 | 20 | 12 | 23 | 28 | 16 |
| 14 | 33 | 7 | 19 | 39 | 17 |
| 18 | 25 | 5 | 20 | 37 | 24 |
| 14 | 33 | 9 | 16 | 43 | 13 |
| 16 | 19 | 14 | 32 | 42 | 16 |
| 10 | 38 | 12 | 24 | 36 | 19 |
| 11 | 26 | 14 | 23 | 35 | 17 |
| 18 | 30 | 11 | 17 | 41 | 19 |

Ordinary One-Way ANOVA $P<0.0001$; Brown-Forsythe test $P<0.0001$; Barlett's test $P<0.0001$

##### Embryonic stage (%) - I359M

| | <20 cells | 20 cells to | post gastrula |
|---|---|---|---|
| **Control (JIP3105)** | 27 | 74 | |
| **I359M (JIP3115)** | 0 | 81 | 20 |
| **kcnl-1(0) (JIP3131)** | 62 | 45 | |

| | | | **Control vs I359M** | **Control vs *kcnl-1 (0)*** | ***kcnl-1 (0)* vs I359M** |
|---|---|---|---|---|---|
| **Fisher Exact + Bonferroni correction** | **< 20 cells** | P value | 0.0000 | 0.0000 | 0.0000 |
| | | Correction | 0.0000 | 0.0000 | 0.0000 |
| | **20 cells to** | P value | 0.3178 | 0.0000 | 0.0000 |
| | | Correction | 0.9533 | 0.0000 | 0.0000 |
| | **post gastrula** | P value | 0.0000 | 1.0000 | 0.0000 |
| | | Correction | 0.0000 | 1.0000 | 0.0000 |

**Figure S2 A**

#### Brood size - I359M

| Control (JIP3105) | I359M (JIP3115) | *kcnl-1(0)* (JIP3131) |
|---|---|---|
| 286 | 108 | 267 |
| 270 | 212 | 305 |
| 256 | 31 | 266 |
| 264 | 186 | 304 |
| 261 | 173 | 239 |
| 302 | 189 | 262 |
| 262 | 157 | 298 |
| 321 | 202 | 292 |
| 240 | 185 | 259 |
| 312 | 142 | 237 |

Kruskal-Wallis test P<0.0001

**Figure S2 A**

#### Embryos in utero - V530L ZLS3 variants (independent lines)

| Control (JIP3105) | V530L (JIP3107) |
|---|---|
| 16 | 30 |
| 15 | 24 |
| 20 | 37 |
| 15 | 31 |
| 20 | 31 |
| 20 | 33 |
| 13 | 33 |
| 11 | 18 |
| 17 | 29 |
| 14 | 17 |
| 14 | 32 |
| 19 | 31 |
| 18 | 30 |
| 16 | 32 |
| 15 | 33 |
| 23 | 35 |
| 21 | 40 |
| 18 | 34 |
| 19 | 21 |
| 19 | 29 |
| 15 | 34 |
| 16 | 31 |
| 16 | 27 |
| 17 | 29 |
| 14 | 31 |
| 14 | 25 |
| 21 | 23 |
| 14 | 35 |
| 10 | 34 |
| 18 | 35 |

Mann Whitney test
P value <0.0001

| Control (JIP3105) | S436C (JIP3109) |
|---|---|
| 10 | 41 |
| 22 | 37 |
| 15 | 39 |
| 13 | 43 |
| 12 | 43 |
| 14 | 42 |
| 14 | 46 |
| 13 | 42 |
| 9 | 46 |
| 9 | 41 |
| 12 | 49 |
| 11 | 44 |
| 11 | 33 |
| 14 | 31 |
| 10 | 31 |
| 22 | 46 |
| 13 | 38 |
| 13 | 34 |
| 19 | 31 |
| 13 | 41 |
| 14 | 43 |
| 12 | 35 |
| 10 | 38 |
| 11 | 40 |
| 13 | 29 |
| 10 | 33 |
| 9 | 35 |
| 12 | 40 |
| 10 | 38 |
| 12 | |

Mann Whitney test
P value <0.0001

**Figure S2 B**

#### Embryonic stage (%) - G350D (independent lines)

| First repetition | | Second repetition | |
|---|---|---|---|
| Control (JIP3105) | G350D (JIP3128) | Control (JIP3105) | G350D (JIP3128) |
| 15 | 14 | 18 | 17 |
| 18 | 14 | 12 | 18 |
| 15 | 20 | 10 | 17 |
| 11 | 12 | 14 | 15 |
| 13 | 12 | 18 | 18 |
| 18 | 12 | 16 | 19 |
| 17 | 19 | 13 | 11 |
| 12 | 16 | 16 | 15 |
| 18 | 11 | 15 | 14 |
| 18 | 11 | 14 | 16 |
| 13 | 12 | 16 | 18 |
| 17 | 15 | 12 | 17 |
| 11 | 13 | 13 | 17 |
| 10 | 14 | 16 | 16 |
| 15 | 15 | 15 | 15 |
| 14 | 17 | 12 | 14 |
| 14 | 11 | 10 | 15 |
| 11 | 13 | 11 | 18 |
| 13 | 12 | 16 | 16 |
| 9 | 11 | 11 | 15 |
| 14 | 12 | 11 | 19 |
| 13 | 14 | 13 | 17 |
| 11 | 11 | 19 | 15 |
| 12 | 12 | 15 | 16 |
| 14 | 13 | 12 | 19 |
| 13 | 11 | 12 | 12 |
| 13 | 16 | 18 | 16 |
| 15 | | 13 | 15 |
| 13 | | 17 | 12 |
| 9 | | 17 | 12 |

Mann Whitney test
P value = 0.1930

| First repetition | | Second repetition | |
|---|---|---|---|
| Control (JIP3105) | G350D (JIP3129) | Control (JIP3105) | G350D (JIP3129) |
| 12 | 17 | 17 | 18 |
| 12 | 20 | 19 | 15 |
| 14 | 17 | 16 | 13 |
| 12 | 15 | 14 | 18 |
| 14 | 14 | 24 | 20 |
| 15 | 19 | 16 | 15 |
| 10 | 15 | 11 | 22 |
| 15 | 14 | 15 | 19 |
| 15 | 14 | 26 | 15 |
| 15 | 21 | 12 | 25 |
| 13 | 24 | 16 | 21 |
| 13 | 17 | 15 | 19 |
| 15 | 13 | 22 | 13 |
| 16 | 16 | 18 | 15 |
| 18 | 32 | 16 | 12 |
| 14 | 17 | 19 | 14 |
| 11 | 13 | 14 | 17 |
| 16 | 19 | 14 | 20 |
| 13 | 17 | 18 | 16 |
| 14 | 15 | 14 | 11 |
| 10 | 19 | 23 | 11 |
| 15 | 12 | 17 | 19 |
| 15 | 14 | 17 | 18 |
| 16 | 10 | 21 | 10 |
| 16 | 16 | 16 | 15 |
| 22 | 12 | 15 | 13 |
| 24 | 19 | 14 | 13 |
| 15 | 14 | 15 | 18 |
| 13 | 10 | 20 | 18 |
| 13 | | | 17 |

Mann Whitney test
P value = 0.3307

#### Figure S2 B

##### Embryonic stage (%) - G350D (independent lines)

| | <20 cells | 20 cells to | post gastrula |
|---|---|---|---|
| **Control (JIP3105)** | 27 | 67 | 0 |
| **G350D (JIP3128) line 2** | 14 | 80 | 0 |

Fisher Exact + Bonferroni

| Category | P value | Correction |
|---|---|---|
| <20 cells | 0.0333 | 0.6650 |
| 20 cells to gastrula | 0.6650 | 0.0665 |

| | <20 cells | 20 cells to | post gastrula |
|---|---|---|---|
| **Control (JIP3105)** | 38 | 62 | 0 |
| **G350D (JIP3129) line 3** | 4 | 110 | 0 |

Fisher Exact + Bonferroni

| Category | P value | Correction |
|---|---|---|
| <20 cells | 0.0000 | 0.0000 |
| 20 cells to gastrula | 0.0000 | 0.0000 |

**Figure S2 C**

#### Embryos in utero - NEDMAB variants (independent lines)

| Control (JIP3105) | G362S (JIP3120) | *kcnl-1(0)* (JIP3131) |
|---|---|---|
| 13 | 10 | 13 |
| 9 | 7 | 10 |
| 18 | 9 | 8 |
| 14 | 12 | 10 |
| 12 | 8 | 13 |
| 10 | 5 | 6 |
| 12 | 10 | 17 |
| 19 | 8 | 10 |
| 12 | 13 | 8 |
| 14 | 10 | 10 |
| 11 | 9 | 19 |
| 14 | 12 | 8 |
| 12 | 10 | 8 |
| 12 | 7 | 10 |
| 10 | 8 | 11 |
| 13 | 11 | 11 |
| 13 | 10 | 8 |
| 10 | 14 | 9 |
| 15 | 15 | 14 |
| 14 | 8 | 8 |
| 14 | 9 | 8 |
| 13 | 10 | 10 |
| 10 | 10 | 13 |
| 11 | 7 | 8 |
| 18 | 9 | 15 |
|  | 10 | 10 |
|  | 5 | 10 |
|  | 8 | 11 |
|  | 9 | 10 |

Kruskal-Wallis test
P value<0.0001

| Control (JIP3105) | L432P (JIP3126) | kcnl-1(0) (JIP3131) |
|---|---|---|
| 11 | 15 | 11 |
| 14 | 14 | 10 |
| 20 | 15 | 11 |
| 12 | 8 | 12 |
| 8 | 12 | 9 |
| 17 | 10 | 17 |
| 15 | 10 | 9 |
| 15 | 6 | 11 |
| 10 | 12 | 8 |
| 15 | 7 | 8 |
| 13 | 13 | 10 |
| 10 | 9 | 6 |
| 18 | 11 | 7 |
| 17 | 8 | 8 |
| 15 | 11 | 8 |
| 16 | 15 | 10 |
| 14 | 15 | 12 |
| 12 | 14 | 8 |
| 15 | 10 | 11 |
| 13 | 11 | 12 |
| 15 | 11 | 21 |
| 14 | 10 | 10 |
| 16 | 12 | 14 |
| 12 | 10 | 14 |
| 10 | 20 | 9 |
| 11 | 13 | 17 |
| 15 | 12 | 13 |
| 10 | 8 | 9 |
| 14 | 4 | 15 |
| 13 | 8 |  |
|  | 10 |  |

Kruskal-Wallis test
P value 0.0007

**Figure S2 C**

#### Embryos in utero - NEDMAB variants (independent lines)

| Control (JIP3105) | L388V (JIP3122) | kcnl-1(0) (JIP3131) |
|---|---|---|
| 19 | 11 | 9 |
| 16 | 15 | 7 |
| 13 | 9 | 11 |
| 15 | 13 | 7 |
| 16 | 9 | 12 |
| 13 | 6 | 12 |
| 12 | 12 | 8 |
| 20 | 14 | 7 |
| 13 | 11 | 9 |
| 11 | 10 | 10 |
| 14 | 11 | 7 |
| 10 | 9 | 11 |
| 15 | 6 | 12 |
| 15 | 13 | 10 |
| 14 | 14 | 9 |
| 15 | 7 | 11 |
| 20 | 6 | 18 |
| 22 | 8 | 4 |
| 24 | 13 | 12 |
| 22 | 9 | 10 |
| 21 | 13 | 9 |
| 15 | 9 | 9 |
| 18 | 14 | 9 |
| 13 | 9 | 7 |
| 18 | 7 | 10 |
| 17 | 8 | 6 |
| 15 | 8 | 8 |
| 16 | 16 | 13 |
| 15 | 12 | 9 |
| 14 | 18 | 11 |

Kruskal-Wallis test
P value<0.0001

| Control (JIP3105) | I288S (JIP3113) | *kcnl-1(0)* (JIP3131) |
|---|---|---|
| 13 | 6 | 11 |
| 16 | 9 | 11 |
| 12 | 11 | 10 |
| 13 | 10 | 4 |
| 14 | 6 | 18 |
| 13 | 16 | 9 |
| 12 | 11 | 13 |
| 13 | 8 | 10 |
| 16 | 7 | 10 |
| 11 | 14 | 12 |
| 14 | 7 | 12 |
| 17 | 16 | 10 |
| 10 | 14 | 8 |
| 13 | 17 | 20 |
| 12 | 14 | 8 |
| 20 | 13 | 9 |
| 16 | 16 | 9 |
| 17 | 9 | 16 |
| 15 | 9 | 15 |
| 14 | 8 | 11 |
| 16 | 11 | 10 |
| 11 | 10 | 10 |
| 14 | 6 | 11 |
| 19 | 9 | 13 |
| 20 | 4 | 11 |
| 14 | 8 | 6 |
| 15 | 13 | 12 |
| 11 | 20 | 15 |
| 16 | 12 | 10 |
| 12 | 6 | 14 |

Kruskal-Wallis test
P value <0.0001

**Figure S2 C**

#### Embryos in utero - NEDMAB variants (independent lines)

| Control (JIP3105) | G371E (JIP3111) | *kcnl-1(0)* (JIP3131) |
|---|---|---|
| 19 | 21 | 18 |
| 14 | 18 | 17 |
| 21 | 16 | 20 |
| 13 | 19 | 16 |
| 19 | 12 | 17 |
| 13 | 16 | 17 |
| 13 | 13 | 16 |
| 24 | 19 | 14 |
| 19 | 18 | 17 |
| 18 | 16 | 20 |
| 18 | 14 | 9 |
| 17 | 15 | 16 |
| 16 | 16 | 19 |
| 23 | 7 | 12 |
| 22 | 13 | 18 |
| 21 | 11 | 10 |
| 12 | 11 | 11 |
| 24 | 17 | 8 |
| 11 | 10 | 12 |
| 16 | 9 | 14 |
| 21 | 9 | 7 |
| 19 | 8 | 18 |
| 22 | 11 | 14 |
| 18 | 12 | 22 |
| 17 | 15 | 20 |
| 16 | 17 | 12 |
| 19 | 13 | 16 |
| 18 | 16 | 19 |
| 21 | 14 | 14 |
| 23 | 14 | 18 |

Ordinary One-Way ANOVA P<0.0001; Brown-Forsythe test P0.5577; Barlett's test P 0.6298

| Control (JIP3105) | I359M (JIP3116) | *kcnl-1(0)* (JIP3131) |
|---|---|---|
| 18 | 26 | 9 |
| 14 | 32 | 18 |
| 13 | 32 | 9 |
| 18 | 42 | 8 |
| 12 | 38 | 11 |
| 18 | 35 | 10 |
| 13 | 29 | 14 |
| 14 | 27 | 11 |
| 17 | 23 | 7 |
| 12 | 35 | 10 |
| 16 | 39 | 8 |
| 15 | 39 | 7 |
| 15 | 36 | 8 |
| 10 | 27 | 7 |
| 18 | 44 | 17 |
| 13 | 40 | 11 |
| 19 | 33 | 10 |
| 21 | 35 | 17 |
| 19 | 24 | 9 |
| 13 | 51 | 11 |
| 15 | 24 | 13 |
| 11 | 29 | 6 |
| 16 | 32 | 9 |
| 15 | 30 | 11 |
| 12 | 33 | 18 |
| 19 | 44 | 9 |
| 12 | 34 | 9 |
| 18 | | 9 |
| 12 | | 11 |
| 12 | | 6 |

Ordinary One-Way ANOVA P<0.0001; Brown-Forsythe test P 0.0003; Barlett's test P<0.0001

#### Figure S3

##### Brood size - NEDMAB variants

| Control (JIP3105) | G362S (JIP3121) | *kcnl-1(0)* (JIP3131) |
|---|---|---|
| 250 | 281 | 305 |
| 122 | 309 | 234 |
| 271 | 279 | 274 |
| 299 | 259 | 147 |
| 254 | 281 | 293 |
| 314 | 289 | 312 |
| 262 | 277 | 253 |
| 268 | 282 | 223 |
| 323 | 246 | 255 |
| 290 | 243 | 309 |

Kruskal-Wallis test P=0.8981

| Control (JIP3105) | L432P (JIP3124) | *kcnl-1(0)* (JIP3131) |
|---|---|---|
| 321 | 298 | 303 |
| 322 | 294 | 298 |
| 175 | 299 | 324 |
| 310 | 293 | 354 |
| 322 | 305 | 265 |
| 312 | 339 | 314 |
| 328 | 350 | 349 |
| 325 | 361 | 256 |
| 332 | 334 | 311 |
| 322 | 284 | 409 |

Kruskal-Wallis test P=0.9251

| Control (JIP3105) | L388V (JIP3123) | *kcnl-1(0)* (JIP3131) |
|---|---|---|
| 252 | | 282 |
| 169 | 222 | 327 |
| 337 | 233 | 270 |
| 231 | 239 | 305 |
| 255 | 321 | 294 |
| 272 | 286 | 301 |
| 289 | 281 | 309 |
| 365 | 295 | 297 |
| | 314 | 346 |
| | 365 | 269 |

Ordinary One-Way ANOVA
P=0.4123; Brown-Forsythe test
P=0.1358; Barlett's test

| Control (JIP3105) | I288S (JIP3114) | kcnl-1(0) (JIP3131) |
|---|---|---|
| 324 | 288 | 183 |
| 258 | 291 | 286 |
| 266 | 336 | 283 |
| 252 | 318 | 282 |
| 227 | 314 | 305 |
| 246 | 304 | 240 |
| 176 | 280 | 231 |
| 256 | 298 | 316 |
| 256 | 285 | 310 |
| 278 | 271 | 321 |

Ordinary One-Way ANOVA
P=0.0307; Brown-Forsythe test
P=0.3582; Barlett's test

| Control (JIP3105) | G371E (JIP3110) | kcnl-1(0) (JIP3131) |
|---|---|---|
| 286 | 315 | 285 |
| 314 | 330 | 318 |
| 271 | 359 | 338 |
| 293 | 357 | 278 |
| 264 | 285 | 320 |
| 285 | 327 | 231 |
| 367 | 298 | 287 |
| 325 | 345 | 367 |
| 301 | 372 | 324 |
| 258 | 290 | 339 |

Ordinary One-Way ANOVA
P=0.1368; Brown-Forsythe test
P=0.8531; Barlett's test

#### Appendix 2

In [7]:

```
# Introduction
# This appendix contains the code to perform Fisher's Exact Test and Bonferroni correction,
# with three examples representing different categories or strains that can be analyzed in the Embryonic Stage
# Specifically, the test for I359M, shown in Figure 4C, is reported here for the version of the code handling 3
# The test for the G350D line 2, shown in Figure S2B, corresponds to the version handling 2 categories / 2 stra
# Finally, the test for V530L, shown in Figure 2C, represents the version handling 3 categories / 3 strains.

import numpy as np
import pandas as pd
import scipy.stats as stats
from statsmodels.stats.multitest import multipletests
import matplotlib.pyplot as plt

#1. INPUT DATA
# Create the 2x3 contingency table
data = {
     'Category': ['<20 cells', '20 cells to gastrula', 'post-gastrula'],
   'Control': [27, 74, 0],  # Data for the Control strain
    'I359M': [0, 81, 20],     # Data for the I359M strain
    'kcnl-1(0)': [62, 45, 0]  # Data for the null strain
}

# Convert to DataFrame
df = pd.DataFrame(data)

#  2. CALCULATE P-VALUES FOR EACH COMPARISON
comparisons = [("Control", "I359M"), ("Control", "kcnl-1(0)"), ("I359M", "kcnl-1(0)")]
results = []

for comp1, comp2 in comparisons:
    p_values = []
    for i in range(len(df)):
    # Create a 2x2 contingency table for the comparison
    # We analyze one category at a time, treating it as "in the category" vs. "not in the category".
    # The table consists of:
    # - The count of samples belonging to the category for each strain.
    # - The count of samples not belonging to that category (sum of the other categories).
        table = np.array([
            [df[comp1][i], df[comp2][i]],
            [df[comp1].sum() - df[comp1][i], df[comp2].sum() - df[comp2][i]]
        ])
        # Perform Fisher's Exact Test
        _, p_value = stats.fisher_exact(table, alternative='two-sided')
        p_values.append(p_value)
    # Append results for this comparison
    results.append((comp1, comp2, p_values))

#  3. MULTIPLE COMPARISONS CORRECTION (BONFERRONI)
adjusted_results = []
for comp1, comp2, p_values in results:
    _, p_adjusted, _, _ = multipletests(p_values, method='bonferroni')
    adjusted_results.append((comp1, comp2, p_values, p_adjusted))

# Print the statistical test results
print("Results of Fisher's Exact Test (with Bonferroni correction):")
for comp1, comp2, p_values, p_adjusted in adjusted_results:
    print(f"\nComparison: {comp1} vs {comp2}")
    for i in range(len(df)):
        print(f"  {df['Category'][i]}: p-value = {p_values[i]:.4f}, corrected p-value = {p_adjusted[i]:.4f}")

#  4. CALCULATE PERCENTAGES FOR THE CHART
percent_data = pd.DataFrame({
    'Category': df['Category'],
    'Control': (df['Control'] / df['Control'].sum()) * 100,
    'I359M': (df['I359M'] / df['I359M'].sum()) * 100,
    'kcnl-1(0)': (df['kcnl-1(0)'] / df['kcnl-1(0)'].sum()) * 100
})

#  5. CREATE STACKED BAR CHART
fig, ax = plt.subplots(figsize=(4, 3))

# Set the colors for the categories
colors = ['#3F56A6', '#75B9DE', '#85C66C']

# Define x-axis labels
x_labels = ['Control', 'I359M', 'kcnl-1(0)']

# Stacked bar plots
for i, category in enumerate(df['Category']):
    bottom_values = np.zeros(len(x_labels)) if i == 0 else percent_data.iloc[:i, 1:].sum().values
```

```
    plt.bar(x_labels, percent_data.iloc[i, 1:], bottom=bottom_values, color=colors[i], label=category)

# Add labels, legend, and title
plt.ylabel('Percentage')
plt.title('Distribution of Cells')
plt.legend(title='Category')

# Add labels and title
ax.set_ylabel('Percentage (%)')
ax.set_title('Distribution of Categories by Strain')
ax.set_ylim(0, 100)  # Set the upper limit of the Y-axis to 100%

# Add the legend
ax.legend(title='Developmental Stages', bbox_to_anchor=(1.05, 1), loc='upper left')

# Rotate X-axis labels at 45 degrees
plt.xticks(rotation=45, ha='right')

# Adjust layout for better readability
plt.tight_layout()

# Save the chart as a PDF file for editing in vector graphic software
plt.savefig("graph.pdf", format='pdf', bbox_inches='tight')

# Show the chart
plt.show()
```

```
Results of Fisher's Exact Test (with Bonferroni correction):

Comparison: Control vs I359M
  <20 cells: p-value = 0.0000, corrected p-value = 0.0000
  20 cells to gastrula: p-value = 0.3178, corrected p-value = 0.9533
  post-gastrula: p-value = 0.0000, corrected p-value = 0.0000

Comparison: Control vs kcnl-1(0)
  <20 cells: p-value = 0.0000, corrected p-value = 0.0000
  20 cells to gastrula: p-value = 0.0000, corrected p-value = 0.0000
  post-gastrula: p-value = 1.0000, corrected p-value = 1.0000

Comparison: I359M vs kcnl-1(0)
  <20 cells: p-value = 0.0000, corrected p-value = 0.0000
  20 cells to gastrula: p-value = 0.0000, corrected p-value = 0.0000
  post-gastrula: p-value = 0.0000, corrected p-value = 0.0000
```

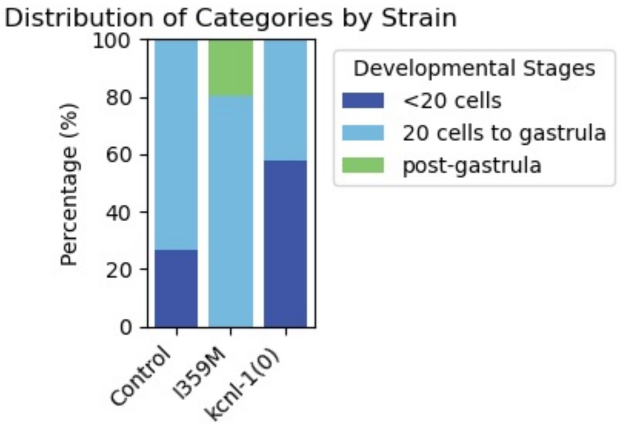

In [ ]:

Loading [MathJax]/jax/output/CommonHTML/fonts/TeX/fontdata.js

In [5]:

```python
# Statistical libraries #
import numpy as np
import pandas as pd
import scipy.stats as stats
from statsmodels.stats.multitest import multipletests
import matplotlib.pyplot as plt

# 1. INPUT DATA
# Define the dataset containing categories and their respective counts for Control and G350D strains.
data = {
    'Category': ['<20 cells', '20 cells to gastrula'],  # Developmental stages
    'Control': [27, 67],  # Observations for the Control strain
    'G350D': [14, 80]     # Observations for the G350D strain
}

# Convert the dictionary into a pandas DataFrame for easier data manipulation
df = pd.DataFrame(data)

# 2. PERFORM FISHER'S EXACT TEST FOR EACH CATEGORY
p_values = []  # List to store p-values
for i in range(len(df)):
    # Create a 2x2 contingency table for Fisher's Exact Test
    # Therefore, we analyze one category at a time, treating it as "in the category" vs. "not in the category".
    # The table consists of:
    # - The count of samples belonging to the category for each strain.
    # - The count of samples not belonging to that category (sum of the other categories).
    table = np.array([
        [df['Control'][i], df['G350D'][i]],  # Samples in the current category for both strains
        [df['Control'].sum() - df['Control'][i], df['G350D'].sum() - df['G350D'][i]]  # Samples in other catego
    ])

    # Perform Fisher's Exact Test (two-sided)
    _, p_value = stats.fisher_exact(table, alternative='two-sided')
    p_values.append(p_value)  # Store the p-value

# 3. MULTIPLE COMPARISONS CORRECTION (BONFERRONI)
# Adjust p-values using the Bonferroni correction to account for multiple comparisons
_, p_adjusted, _, _ = multipletests(p_values, method='bonferroni')

# Print the statistical test results
print("Results of Fisher's Exact Test (with Bonferroni correction):")
for i in range(len(df)):
    print(f"{df['Category'][i]}: p-value = {p_values[i]:.4f}, corrected p-value = {p_adjusted[i]:.4f}")

# 4. CALCULATE PERCENTAGES FOR THE CHART
# Convert absolute counts into percentages to visualize the proportion of each category

df['Control_percent'] = (df['Control'] / df['Control'].sum()) * 100

df['G350D_percent'] = (df['G350D'] / df['G350D'].sum()) * 100

# 5. CREATE STACKED BAR CHART
# Initialize a figure with specified dimensions
fig, ax = plt.subplots(figsize=(4, 3))

# Define colors for each category in the stacked bar chart
colors = ['#3F56A6', '#75B9DE']

# Create stacked bars for 'Control' group
top_control = ax.bar('Control', df['Control_percent'][0], color=colors[0], label=df['Category'][0])
bottom_control = ax.bar('Control', df['Control_percent'][1], bottom=df['Control_percent'][0], color=colors[1], 

# Create stacked bars for 'G350D' group
top_g350d = ax.bar('G350D', df['G350D_percent'][0], color=colors[0])
bottom_g350d = ax.bar('G350D', df['G350D_percent'][1], bottom=df['G350D_percent'][0], color=colors[1])

# Set labels and title
ax.set_ylabel('Percentage (%)')
ax.set_xlabel('Strain')
ax.set_title('Distribution of Categories by Strain (Control vs G350D)')
ax.set_ylim(0, 100)  # Set Y-axis limits to 100% to represent proportions correctly

# Add a legend to distinguish categories
ax.legend(title='Developmental Stages', bbox_to_anchor=(1.05, 1), loc='upper left')

# 6. SAVE THE CHART AS A PDF FILE
# Save the chart for further editing in vector graphic software
plt.savefig("graph.pdf", format='pdf', bbox_inches='tight')

# Show the chart
plt.tight_layout()
plt.show()
```

```
Results of Fisher's Exact Test (with Bonferroni correction):
<20 cells: p-value = 0.0333, corrected p-value = 0.0665
20 cells to gastrula: p-value = 0.0333, corrected p-value = 0.0665
```

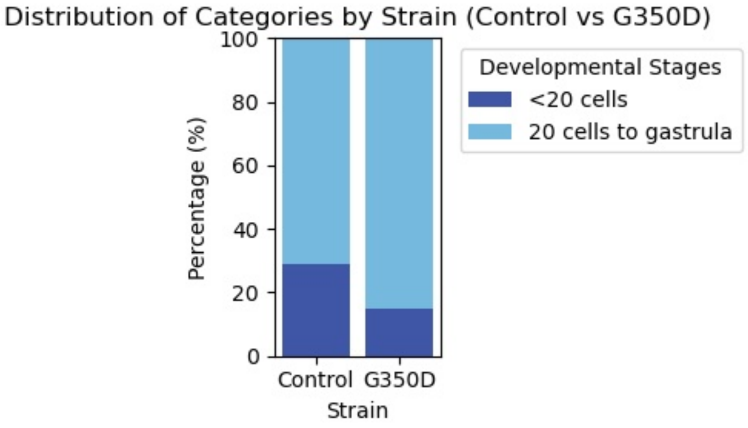

In [ ]:

Loading [MathJax]/jax/output/CommonHTML/fonts/TeX/fontdata.js

In [1]:

```
# Statistical libraries #
import numpy as np
import pandas as pd
import scipy.stats as stats
from statsmodels.stats.multitest import multipletests
import matplotlib.pyplot as plt

# 1. INPUT DATA
# Create the dataset containing categories and their respective counts for Control and V530L strains.
data = {
    'Category': ['<20 cells', '20 cells to gastrula', 'post-gastrula'],  # Developmental stages
    'Control': [34, 68, 0],  # Observations for the Control strain
    'V530L': [0, 78, 25]     # Observations for the V530L strain
}

# Convert the dictionary into a pandas DataFrame
df = pd.DataFrame(data)

# 2. PERFORM FISHER'S EXACT TEST FOR EACH CATEGORY
p_values = []  # List to store p-values
for i in range(len(df)):
    # Create a 2x2 contingency table for Fisher's Exact Test (Even though we have 3 categories, Fisher's Exact 
    # Therefore, we analyze one category at a time, treating it as "in the category" vs. "not in the category".
    # The table consists of:
    # - The count of samples belonging to the category for each strain.
    # - The count of samples not belonging to that category (sum of the other categories).
    table = np.array([
        [df['Control'][i], df['V530L'][i]],  # Samples in the current category for both strains
        [df['Control'].sum() - df['Control'][i], df['V530L'].sum() - df['V530L'][i]]  # Samples in other catego
    # Perform Fisher's Exact Test (two-sided)
    _, p_value = stats.fisher_exact(table, alternative='two-sided')
    p_values.append(p_value)  # Store the p-value

# Adjust the p-values using Bonferroni correction to account for multiple comparisons
_, p_adjusted, _, _ = multipletests(p_values, method='bonferroni')

# Print the statistical test results
print("Results of Fisher's Exact Test (with Bonferroni correction):")
for i in range(len(df)):
    print(f"{df['Category'][i]}: p-value = {p_values[i]:.4f}, corrected p-value = {p_adjusted[i]:.4f}")

# 3. CREATE STACKED BAR CHART
# Convert absolute counts into percentages for visualization

df['Control_percent'] = (df['Control'] / df['Control'].sum()) * 100

df['V530L_percent'] = (df['V530L'] / df['V530L'].sum()) * 100

# Create the chart
fig, ax = plt.subplots(figsize=(4, 3))

# Define colors for each category in the stacked bar chart
colors = ['#3F56A6', '#75B9DE']

# Create stacked bars for 'Control' group
top_control = ax.bar('Control', df['Control_percent'][0], color=colors[0], label=df['Category'][0])
bottom_control_1 = ax.bar('Control', df['Control_percent'][1], bottom=df['Control_percent'][0], color=colors[1]
bottom_control_2 = ax.bar('Control', df['Control_percent'][2], bottom=df['Control_percent'][0] + df['Control_pe

# Create stacked bars for 'V530L' group
top_v530l = ax.bar('V530L', df['V530L_percent'][0], color=colors[0])
bottom_v530l_1 = ax.bar('V530L', df['V530L_percent'][1], bottom=df['V530L_percent'][0], color=colors[1])
bottom_v530l_2 = ax.bar('V530L', df['V530L_percent'][2], bottom=df['V530L_percent'][0] + df['V530L_percent'][1]

# Set labels and title
ax.set_ylabel('Percentage (%)')
ax.set_xlabel('Strain')
ax.set_title('Distribution of Developmental Stages by Strain (Control vs V530L)')
ax.set_ylim(0, 100)  # Set Y-axis limits to 100% to represent proportions correctly

# Add a legend to distinguish categories
ax.legend(title='Developmental Stages', bbox_to_anchor=(1.05, 1), loc='upper left')

# === 4. SAVE THE CHART AS A PDF FILE ===
# Save the chart for further editing in vector graphic software
plt.savefig("graph.pdf", format='pdf', bbox_inches='tight')

# Show the chart
plt.tight_layout()
plt.show()
```

```
Results of Fisher's Exact Test (with Bonferroni correction):
<20 cells: p-value = 0.0333, corrected p-value = 0.0665
20 cells to gastrula: p-value = 0.0333, corrected p-value = 0.0665
```

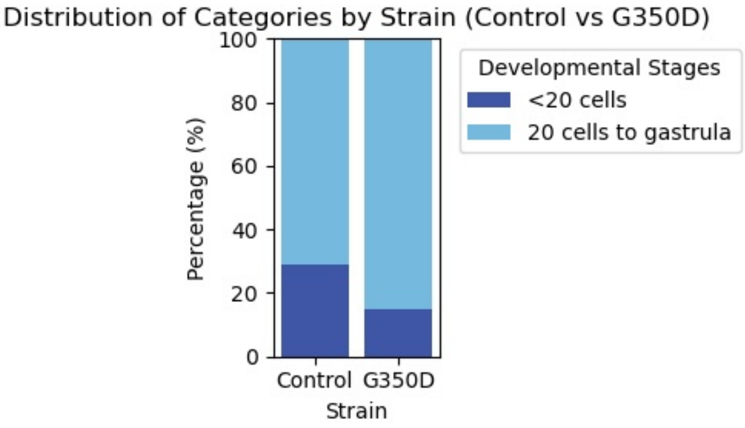

In [ ]:

Loading [MathJax]/jax/output/CommonHTML/fonts/TeX/fontdata.js
